# BiNVaST: niche boundary-preserving and synthetic data-guided variational autoencoders for concordant inference across high-resolution spatial transcriptomic platforms

**DOI:** 10.64898/2026.10.15.759369

**Authors:** Jingxuan Zhang, Lei Li, Limeng Liu, Brooke Fridley, Cheng Cheng, Jasmine Plummer, Gang Wu, Jason Chiang, Qian Li

## Abstract

High-resolution spatial transcriptomics (ST) enables subcellular spatially resolved transcript profiling but remains difficult to annotate in complex tissues because of gene coverage and super sparseness in expression. These challenges are amplified in cancer research, where neoplastic cell states and rare immune cells in tumor microenvironment are often poorly captured by generic references and/or cell typing tools exclusively built on molecular profile. Here, we present BiNVaST, a niche boundary-preserving and dual-branch variational autoencoder fused with statistical simulator for reliable cell typing across multiple spatial platforms. It generates platform-adjusted in silico training data by integrating query spatial profile and user-defined pre-annotated single-cell reference through Singular Value Decomposition and Optimal Transport, combining molecular similarity and spatial proximity to predict segmented cell labels. BiNVaST was benchmarked against existing methods across Xenium, CosMx, Stereo-seq, Visium HD platforms using six specimens from four tissue types. This tool is sensitive in detecting divergent tumor cell states and improves the cross-platform concordance, distinguishing disease-specific neoplastic and non-neoplastic cellular subpopulations and supporting robust cellular or niche inferences in heterogeneous tissues.

## Background

Low-resolution spatial transcriptomics (ST) platforms, such as 10x Visium and earlier array-based methods, measure gene expression from spots that contain a mixture of *>* 10 cells. Hence, cell type deconvolution is essential for estimating the cellular composition of each spot. Most approaches use single-cell or single-nucleus RNA-seq data as a reference and model spot-level gene expression as weighted sum of cell-type-specific expression, similar to bulk deconvolution, such as RCTD [1], cell2location [2], SPOTlight[3], stereoscope [4], and Tangram [5]. These methods have enabled spatial analysis of complex tissues, including tumors, normal brain, and immune microenvironments, but their accuracy depends strongly on reference quality, marker specificity, sequencing depth, spot size, and the similarity between single-cell references and the target (query) spatial tissues.

Recent advancement in high-resolution ST has shifted cell labeling paradigm from mixture-spot deconvolution toward cell-aware or molecule-aware annotation. For sequencing-based assays such as Visium HD, the raw data are captured in small spatial bins rather than pre-defined cells, leading to extremely low counts and severe zero-inflation in gene expression. Hence, Visium HD cell labeling may require histology-derived nuclei/cell segmentation, followed by aggregation of bins to cells that can increase transcript abundance. On the other hand, imaging-based platforms such as MERFISH, CosMx, Xenium, Atera detected individual RNA molecules with subcellular coordinates, followed by transcript-to-cell assignment. Vendor pipelines provide baseline segmentation [6], while research tools such as Baysor [7] can improve cell calling by jointly modeling molecule positions, gene identity, and local spatial density. Other tools such as Cellpose [8], StarDist [9], and Mesmer[10] are widely used for image-based nuclei or whole-cell segmentation, and newer workflows such as Bin2cell [11] and STHD [12] were designed for Visium HD binning of expression data.

For both sequencing- and imaging-based platforms, the post-segmentation cell phenotyping is usually performed by transferring annotations from tissue-type-matched single-cell RNA-seq references with tools such as SingleR [13], Seurat[14], or a probabilistic mapping approaches like STHD. These tools differ in whether they assign one dominant label per cell/bin, estimate mixtures of multiple cell types, or map single-cell reference states into tissue space. In practice, high-resolution platforms reduce but do not eliminate ambiguity: (1) targeted panels in Xenium/CosMx/MERFISH can limit marker coverage; (2) segmented cells in Visium HD may still capture partial cells or extracellular RNA; (3) tumor tissues often contain transcriptional continua that do not fit clean and discrete cell states. Thus, the best practice is to combine morphology-aware segmentation, reference-guided annotation, marker-gene validation, and spatial neighborhood consistency rather than relying on a single automated label.

A major challenge in high-resolution ST cell typing arises from disease- or tissue-specific heterogeneity. Multiple cutting-edge platforms, e.g., Xenium V1, Xenium Prime 5K panels, CosMx, Stereo-seq, Visium HD and emerging platforms such as Atera, a more advanced platform in future environments, are favorable for complex tissues, including but not limited to rare tumors. However, inferring tumor cell lineages and/or tumor microenvironment (TME) with these technologies remains extremely challenging due to ambiguous neoplastic cell evolution, immune microenvironment, and therapy-induced immune-evasive niches. A recent study [15] found that CosMx 6K detected a higher total number of transcripts but exhibited reduced spatial variation compared to Xenium 5K, while the annotated clusters in each ST platform (Visium HD, CosMx 6K, Xenium 5K) were poorly aligned with the orthogonal PhenoCycler Fusion spatial clusters in the tissue-matched adjacent sections (cluster proportions Pearson correlation *<* 0.5). Popular cell phenotyping tools like Seurat[14], SingleR[13], SELINA[16], Celltypist[17], TACCO[18] showed stable performance in normal cell annotation, but they exposed discordance in distinguishing divergent tumor cell states and rare immune cell subpopulations due to ambiguous cell markers. The agreement across these methods also varies between platforms, i.e., Xenium 5K showing higher concordance. Moreover, a new deep learning tool Azimuth, recently developed for (Visium HD) segmented cells, does not support disease-specific or user-defined tumor cell state labeling and only maps malignant cells to similar normal cells in their pre-built non-cancer pan-human single cell reference. Further, the cellular and niche annotations and inference for Visium HD profiles using 2×2 um bins vs. segmented cells have not been systematically compared.

To resolve these caveats, we develop a dual-branch framework, BiNVaST, with niche <u>B</u>oundary-preserving and sy<u>N</u>thetic data-guided <u>V</u>ariational <u>a</u>utoencoders for high-resolution <u>ST</u> cell typing. This tool employs a novel built-in probabilistic simulator to generate platform- and tissue-adjusted synthetic high-resolution ST data for training, enabling cross-platform-concordant spot annotation using disease-specific and user-defined reference scRNA-seq. We rigorously benchmarked the discordance/concordance between competing methods across different ST panels or platforms, based on neoplastic and non-neoplastic cell labels relative to H&E histologic and/or immunohistochemical (IHC) ground truth.

In the benchmarking study, we evaluated each tool for the concordance across spatial platforms by (1) overall cellular abundance with H&E validation, (2) cell-type-specific marker gene inference. The performance of competing methods will be demonstrated by six specimens across four tissue types, i.e., human normal kidney, ovarian cancer (OV), colorectal cancer (CRC), pediatric high grade glioma (pHGG), profiled by imaging-based (Xenium, CosMx) and sequencing-based (Visium HD, Stereo-seq) platforms. Notably, BiNVaST outperformed STHD, Pan-Human Azimuth neural network, Seurat, and SingleR in rigorous cross-resolution, cross-panel, cross-platform evaluations, using pre- and post-cell-segmentation profiles from a Visium HD sample and the adjacent tissue sections profiled by distinct panels or platforms. Our tool exhibited lower variance across platform pairs, cleaner niche boundaries, better recovery of rare tumor states, and supported custom single cell references. Further, BiNVaST accurately annotated TME populations (e.g., endothelial cells, myeloid cells, oligodendrocytes) and distinguished ambiguous neurodevelopmental states of neoplastic cells in pHGG tumor specimen profiled by Xenium 5K. The niches built from BiNVaST predicted cell labels aligned with anatomic structures of cerebral cortex.

## Results

### Overview of BiNVaST

We build a dual-branch variational autoencoders (VAEs) framework that predicts the probabilities of spot-level cell labels in a query tissue section using its high-resolution ST molecular profile and spatial coordinates along with a pre-annotated tissue-type-matched scRNA-seq reference (Figure 1). The input genes are those shared between single cell reference and the query spatial profile, pre-selected by the Variance Stabilizing Transformation [14] in the query sample. Another option is to select the cell-type-informative genes from single cell reference, although this approach usually requires high-quality pre-annotated reference with controlled difference between platforms. Users can manually adjust the input gene list to optimize cell label prediction.

**Fig. 1.**
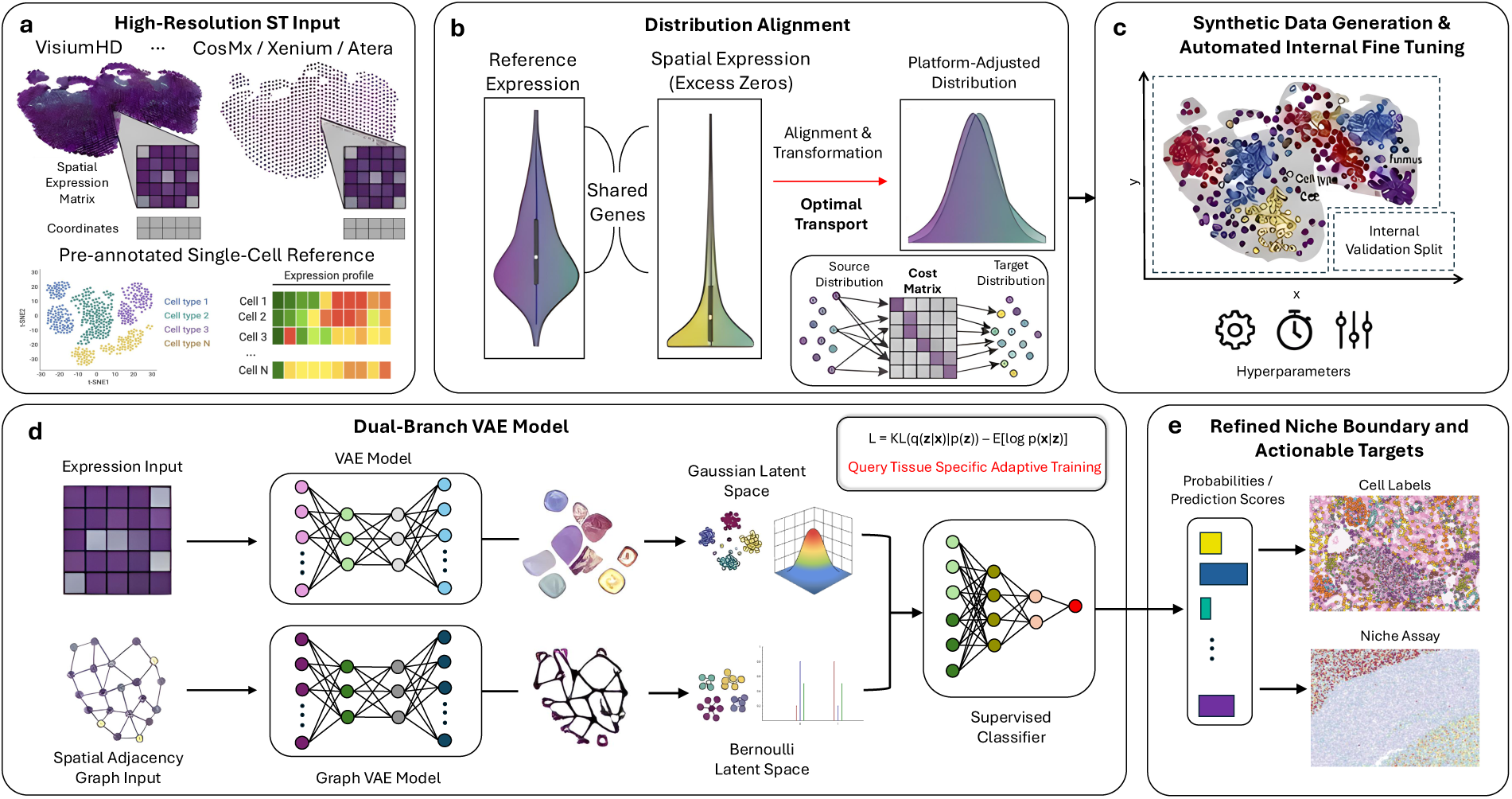
BiNVaST : a framework of generative AI and Statistical simulator. **(a)** Input data include raw expression at single cell resolution and physical x,y coordinates per cell in the query sample, along with pre-annotated scRNA-seq reference expression. **(b)** Generate spatially resolved synthetic expression with pseudo spatial coordinates, adapted to the query sampel profiling platform. **(c)** Fine-tune epoch and batch size hyperparameters by internal validation on synthetic data. **(d)** A dual-brach VAE model predicting cell label probability scores. **(e)** The cell labels can refine domain boundaries by building niche assay.

To train a dual-branch VAE model, we devise a novel spatial simulator that narrows the gap between profiling platforms (i.e., single cell or single nuclei RNA-seq vs. high-resolution ST with cell segmentation), as well as integrates the molecular and cellular organizations in both reference and query tissues. First, we generated synthetic scRNA-seq data from the reference by random sampling with replacement or Generalized Additive Model (GAM) with tensor predictors, and then adjusted the expression distribution by injecting zero-inflation estimated from the query ST data. Next, the synthetic vs. query expressions are further aligned via Singular Value Decomposition (SVD) projection of the query data *Y_q_* onto the basis of reference data 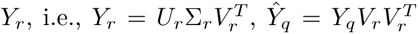. BiNVaST also enables reverse projection of the synthetic data onto the basis of query molecular space as an alternative approach to optimize performance across different technologies. With the platform-aligned expression matrices, we employ Euclidean-cost entropy-regularized Optimal Transport (OT) with a Gaussian (RBF) kernel to map each synthetic cell to a spot in the query tissue, assigning biologically meaningful spatial coordinates to the synthetic molecular profile. This dual-branch VAEs will be pre-trained on the synthetic tissue and then adapted to the query ST profile by re-optimizing the loss function.

The first VAE in this tool is applied to the molecular profile (MVAE), reducing the dimension of input genes via a multi-layer neural network and projecting the low-dimensional embedding to the multivariate Gaussian space. The second branch is a graph-only VAE (GVAE) that takes an adjacency matrix built by KNN from spatial coordinates as input, projecting the latent embedding to a Bernoulli space and reconstructing the neighborhood structure. This graph-based probabilistic model integrates the spots within a proximity defined by their molecular similarity and physical distances, effectively grouping subcellular transcript measurements and distinguishing the cellular subpopulations that can refine niche boundaries. The decoder reconstruction likelihood functions for molecular space 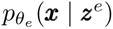 and neighborhood *p_θs_*(***M*** *| **Z**^s^*) from the latent embeddings are formulated as

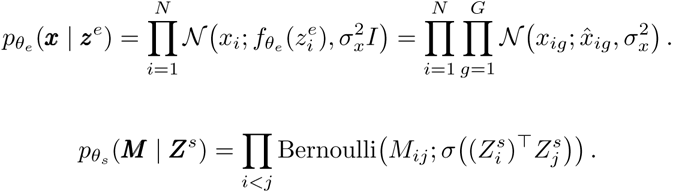

whereas *x_i_* is input expression profile for cell 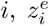 is the reduced-dimension expression latent variables with fixed variance 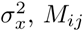 is the edge for spots (cells) *i, j* in the spatial adjacency graph matrix ***M***, and ***Z****^s^* is the spatial latent variables. These two latent representations are then combined to infer cell-type probabilities and improve spatially resolved annotation of cellular niches.

Last, we train a supervised classifier that fuses the Gaussian and Bernoulli latent spaces to predict the probability scores of all pre-defined cell types for each spot, optimizing a supervised cross-entropy loss jointly with the latent-space reconstruction objectives. Cell labels are assigned by the maximum predicted probability when the confidence exceeds a pre-specified threshold (e.g., *p >* 0.6); spots below this threshold are treated as ambiguous, while the learned synthetic and real molecular embeddings are used to assess alignment and support downstream visualization of cellular subpopulations.

### Human Normal Kidney: Visium HD before and after cell segmentation

We conducted a benchmarking study on the Visium HD profile of human normal kidney section at different resolutions: 2×2 *µ*m bin (i.e., before cell segmentation) vs. single cell (i.e., after cell segmentation). STHD was applied to 7,397,943 2 *µ*m bins, using pre-annoated scRNA-seq reference (Figure 2a), while Azimuth (Figure 2b) and BiNVaST (Figure 2c) were applied to 148,055 segmented cells in this Visium HD sample. Each 2*µ*m bin center was transformed into the full-resolution image coordinate system and assigned to a cell if its centroid was within a cell segmentation polygon. Principal cells (PC) and connecting tubule (CNT) were grouped as Cortical Collecting Duct (CCD)-like; Undefined reference cell type (U1, U2) and ambiguous calls by STHD or BiNVaST were treated as Unassigned. In Azimuth predictions, cell labels not supported by organ or predicted as NA were grouped as Unassigned and excluded from agreement denominators.

**Fig. 2.**
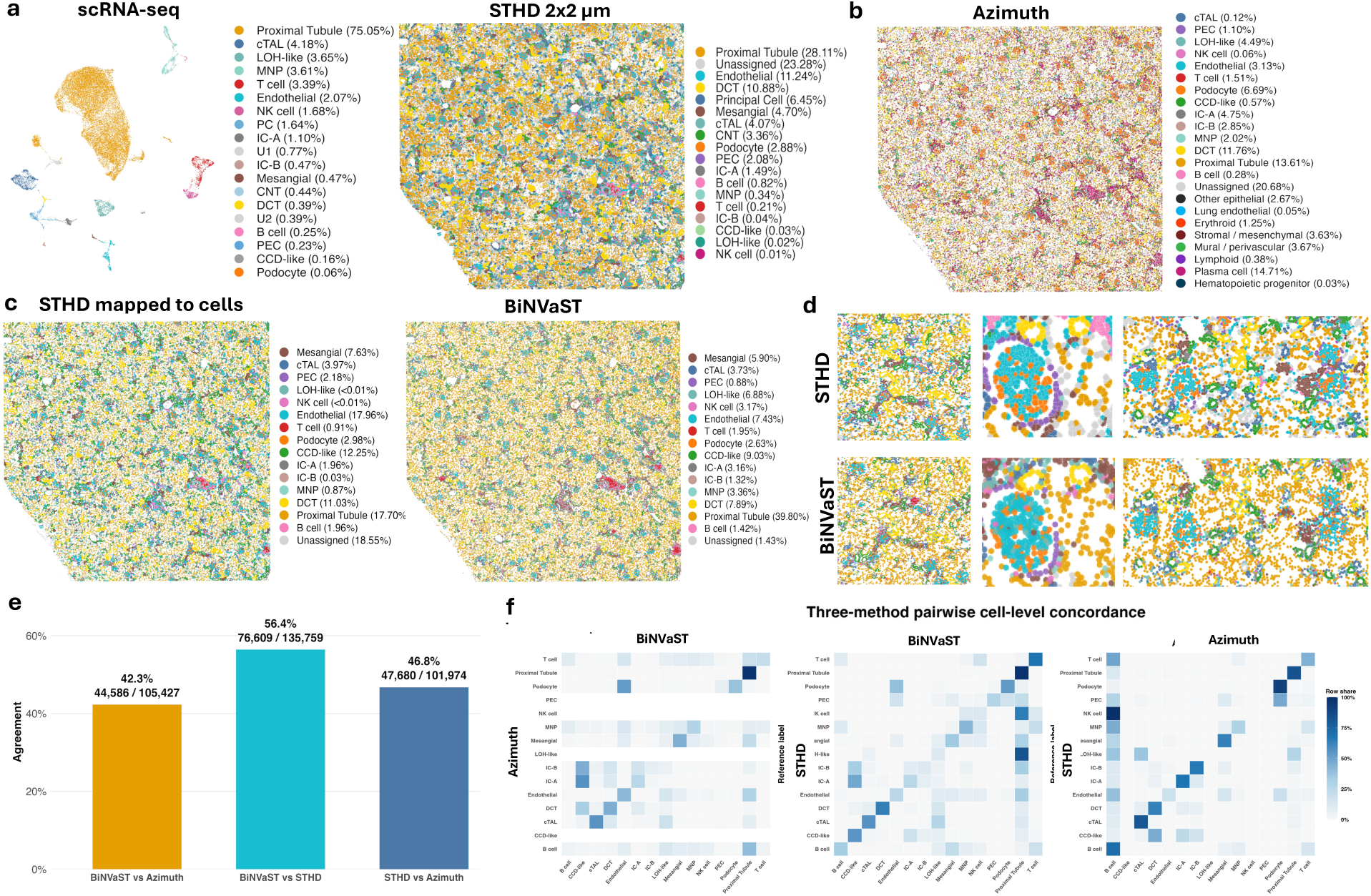
Human normal kidney tissue profiled by Visium HD. **(a)** UMAP for an external normal kidney scRNA-seq reference and spatial map for STHD labels at 2×2 micrometer (*µ*m) resolution without cell segmentation. Cell-type proportions are shown in the legends. DCT: Distal convoluted tubule, CNT: Connecting tubule, LOH: Loop of Henle, cTAL: Cortical thick ascending limb, CCD: Cortical collecting duct, PC: Principal cell, IC: Intercalated cell, PEC: Parietal epithelial cell, NK: Natural Killer, MNP: Mononuclear phagocyte. **(b)** Pan-human Azimuth cell annotation post cell segmentation. **(c)** STHD spot labels mapped to segmented cells (left) and BiNVaST cell-type predictions (right), with cell-type proportions shown for each result. **(d)** Three representative regions comparing local STHD and BiNVaST annotations using the same color key. **(e)**Pairwise cell-level agreement between BiNVaST, Azimuth, and STHD. Labels above the bars report the agreement percentage and the number of concordantly labeled cells over the total number compared. **(f)** Pairwise cell-level confusion matrices for BiNVaST versus Azimuth, BiNVaST versus STHD, and Azimuth versus STHD.

At the bin level, 1,319,428 of 2,169,428 eligible 2*µ*m bins agreed with the Azimuth label of the containing cell (60.8%). At the cell level, the majority STHD label agreed with Azimuth in 34,099 of 61,738 eligible cells (55.2%; Cohen’s *κ* = 0.504).This is a cross-resolution concordance analysis, not a one-to-one comparison of identical units. STHD predicts labels for 2*µ*m bins, whereas Azimuth predicts one label per segmented cell. The cell-majority statistic is therefore the primary comparison; the spot-level statistic is more sensitive to within-cell mixtures and boundary effects. BiNVaST was trained with the target-PCA nearest-cell Gaussian strategy using 2,000 highly variable genes. Proximal Tubule was predicted as the predominant class (39.80%), followed by CCD-like (9.03%), DCT (7.89%), Endothelial (7.43%), LOH-like (6.88%), and Mesangial (5.90%) cells; only 1.43% of cells were labeled Unassigned. The whole-section map showed broad proximal-tubule coverage with spatially localized distal-nephron, vascular, glomerular, stromal, and immune-cell domains. In the common zoom region, BiNVaST reproduced much of the broad spatial organization observed after aggregating STHD 2 *µ*m labels to segmented cells, whereas Azimuth displayed a more heterogeneous local composition and a larger visually unassigned component. It’s worth noting that Pan-Human Azimuth does not allow users to select or load tissue-specific single cell reference. Cell types from other organs or tissues shown in the predicted labels illustrates the limitation of this tool. After excluding Unassigned labels for each method, STHD and Azimuth agreed for 47,680 of 101,974 eligible cells (46.8%; Cohen’s *κ* = 0.415), BiNVaST and Azimuth agreed for 44,586 of 105,427 cells (42.3%; *κ* = 0.360), while BiNVaST and STHD agreed for 76,609 of 135,759 cells (56.4%; *κ* = 0.491; Figure 2e). At the native STHD resolution, agreement was 52.6% between STHD and Azimuth and 63.6% between STHD and BiNVaST. Among 101,011 cells carrying valid labels from all three methods, 35,820 (35.5%) received the same label from every method, while BiNVaST agreed with at least one comparator for 66,074 cells (65.4%). Class-specific comparison against Azimuth was strongest for Proximal Tubule (97.9%), followed by cTAL (56.7%), DCT (45.8%), Mesangial (45.6%), and Endothelial (41.8%); the confusion matrices and per-class results indicate that the remaining disagreement was concentrated among related epithelial states and lower-frequency immune or intercalated-cell classes. The cell-level agreement between three methods are summarized in Figure 2f, showing larger variation Azimuth vs. STHD agreement compared to BiNVaST vs. STHD.

### Ovarian Cancer: Xenium V1, Xenium Prime 5K

We validated BiNVaST against Seurat and SingleR on the Xenium 5K data profiled from an Ovarian Cancer (OV) tissue, where the tissue-matched scRNA-seq is available (Figure 3a (left)) and serves as reference required by each method, as well as provides baseline reference for cell proportions and marker genes. Two primary malignant cell states, i.e., (Non-Proliferative) Tumor Cells and Proliferative Tumor Cells were identified in scRNA-seq, followed by VEGFA+ Tumor Cells and MT-High and Jun+/Fos+ Tumor Cells (Figure 3a (right)). The Inflammatory Tumor Cells were rare in this tissue, identified as a small cluster in scRNA-seq Uniform Manifold Approximation and Projection (UMAP,Figure 3a). Notably, BiNVaST annotated the primary and secondary (VEGFA+, MT-High and Jun+/Fos+) malignant cell states with abundance comparable to the internal scRNA-seq, while accurately identifying the Inflammatory Tumor Cells in a small niche. Seurat and SingleR failed to annotate the major tumor cell states and predicted most malignant cells as VEGFA+ and MT-High and Jun+/Fos+ (Supplementary Figure S1). In the first selected region of interest (ROI), three methods correctly and concordantly annotated endothelial cells (Figure 3c). But the niche of Pericytes, Smooth Muscle cells, and Macrophages were better defined by BiNVaST with only few malignant cells. Seurat and SingleR either labeled more malignant cells or underestimated Pericytes in this ROI (Supplementary Figure S1). A second ROI is the lining of fallopian tube epithelium, the origin site that high grade serous ovarian tumor cell starts [19]. Interestingly, Seurat and SingleR labeled this region as normal epithelial cells, while BiNVaST predicted this region dominated by the major tumor cells pre-defined in the tissue-matched scRNA-seq.

**Fig. 3.**
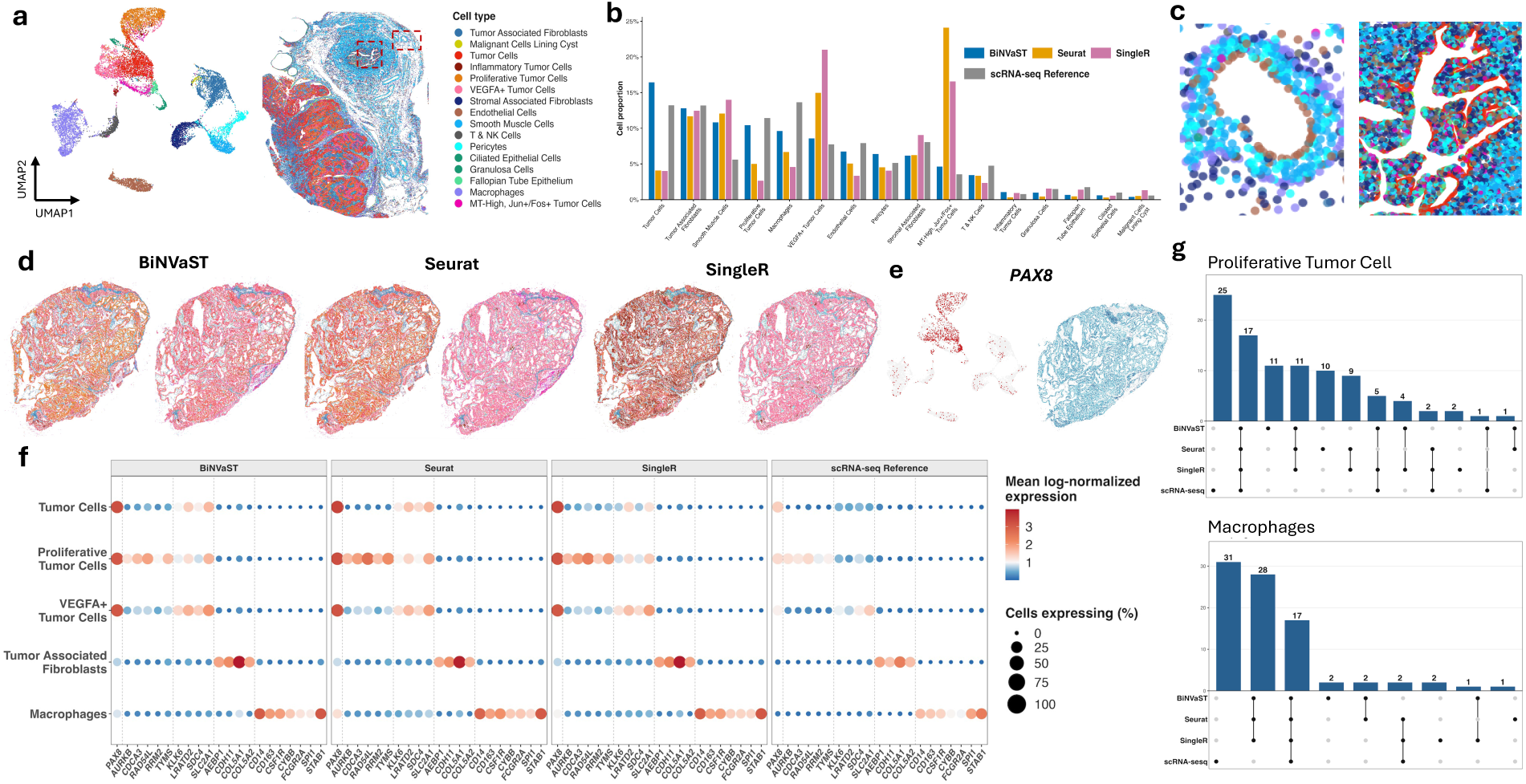
**(a)** UMAP of the ovarian cancer scRNA-seq reference and spatial distribution of the corresponding cell types in the ovarian Xenium 5K tissue. Colors denote the shared cell-type annotation scheme. **(b)** Cell-type proportions estimated by BiNVaST, Seurat label transfer, SingleR with an aggregated reference, and the scRNA-seq reference. **(c)** Representative ROIs shown in panel (a), illustrating predicted cell identities and local cellular organization. **(d)** Cell label spatial maps for OV adjacent sections profiled by Xenium V1 (left) vs. Xenium 5K (right), annotated by BiNVaST, Seurat, and SingleR. The same color key is used across methods and samples. **(e)** *PAX8* expression in the scRNA-seq reference UMAP (left) and OV Xenium 5K (right). **(f)** Bubble plot comparing selected marker-gene expression across BiNVaST, Seurat, SingleR, and the scRNA-seq reference. Rows represent tumor and stromal/immune cell states; dot size indicates the percentage of cells expressing each gene, and color indicates mean log-normalized expression. **(g)** UpSet plots showing intersections among the top marker genes identified by BiNVaST, Seurat, SingleR, and the scRNA-seq reference for proliferative tumor cells (top) and macrophages (bottom).

We conducted a second benchmarking experiment on two adjacent sections of another OV tumor specimens profiled by Xenium V1 (477-gene) panel and Xenium Prime 5K panel, respectively to evaluate the impact of gene panel dimension on cell annotation accuracy. The above scRNA-seq sample was used as an external reference for each tool. In the lower-dimension gene panel, BiNVaST was partially concordant with Seurat (45.03%) and less concordant with SingleR (28.85%), with cell-type-specific agreement visualized in Supplementary Figures xx. In BiNVaST result, 59.07% tumor cells were consistent with Seurat labels. SingleR predicted 42.83% Inflammatory Tumor Cells, 15.76% Proliferative and 4.99% Non-Proliferative Tumor Cells. On the other hand, the cell annotations for an adjacent section in this tumor profiled by Xenium Prime 5K exhibited striking discordance with Xenium V1 panel. Surprisingly, VEGFA+ tumor cells became the dominant malignant cell state in the predictions by Seurat and SingleR, where the major tumor cell states (either proliferative or non-proliferative) were substantially reduced and invisible in the spatial map (Figure 3). On the other hand, using the expanded 5000-gene panel, BiNVaST identified more VEGFA+ Tumor Cells in this tissue while keeping proliferative and non-proliferative cancer cells at reasonable abundance. Tummor Associated Fibroblast was concordantly labeled across two Xenium panels by BiNVaST and Seurat, but SingleR labeled some of these cells as Pericytes. Seurat barely annotated Stromal Associated Fibroblast and did not identify any Malignant Cells Lining Cyst.

To assess the impact of cell labels on downstream inference, we compared marker gene expression between the scRNA-seq vs. Xenium 5K annotated by BiNVaST, Seurat, and SingleR, individually. Differential expression analysis identified reproducible signatures across methods in 5 tumor cell states and 3 immune cell subpopulations, although the malignant cell states labeled by Seurat and SingleR were accurate based on OV tumor cell marker gene *PAX8* (Figure 3e), which was consistently enriched in tumor cells across Xenium and scRNA-seq assay. Proliferative Tumor Cells consistently expressed cell-cycle markers, including *AURKB*, *CDCA3*, and *RAD54L*, whereas Tumor Associated Fibroblasts shared extracellular-matrix and stromal markers such as *AEBP1, CDH11, COL5A1,* and *COL5A2*. Macrophages exhibited the strongest cross-method concordance, with shared expression of *CD14, CD163, CSF1R, CYBB, FCGR2A, ITGAX, SPI1,* and *STAB1*. These results demonstrate that the main tumor cells, stromal and myeloid populations identified in the Xenium 5K data retained transcriptional programs observed in the scRNA-seq reference, supporting the biological validity of the spatial cell-type annotations. Overall, Macrophage identity is the most reproducible across platforms and annotation methods. Tumor-associated fibroblasts also show strong agreement, while Proliferative and Inflammatory tumor states have coherent cell-cycle and interferon-response overlap.

### Colorectal Cancer: Xenium V1, Xenium 5K, CosMx 6K, Stereo-seq

We expanded the concordance benchmarking to different high-resolution ST platforms (i.e., Xenium CosMx, Stereo-seq) in Colorectal Cancer (CRC). The scRNA-seq data obtained from *n* = 8 CRC tumor and normal adjacent tissues with 226, 856 cells in 38 pre-defined cellular subpopulations was used as full single cell reference, whereas one tumor sample is matched to the Xenium V1 (422-gene) sample. We also reduced the full single cell reference to *n* = 16, 564 cells with balanced cell proportions as a secondary reference for benchmarking experiments. With the Xenium V1 panel and full reference, BiNVaST and Seurat clearly defined the niches enriched by macrophages, CD4+ T cells, and neutrophil, while SingleR did not distinguish these cell types, illustrated by ambiguous niche boundaries (Figure 4b). The cell abundance barplot for Xenium and tissue-matched scRNA-seq illustrated the agreement between BiNVaST and Seurat in leading cell populations and their outperformance over SingleR (e.g., Tumor II).

**Fig. 4.**
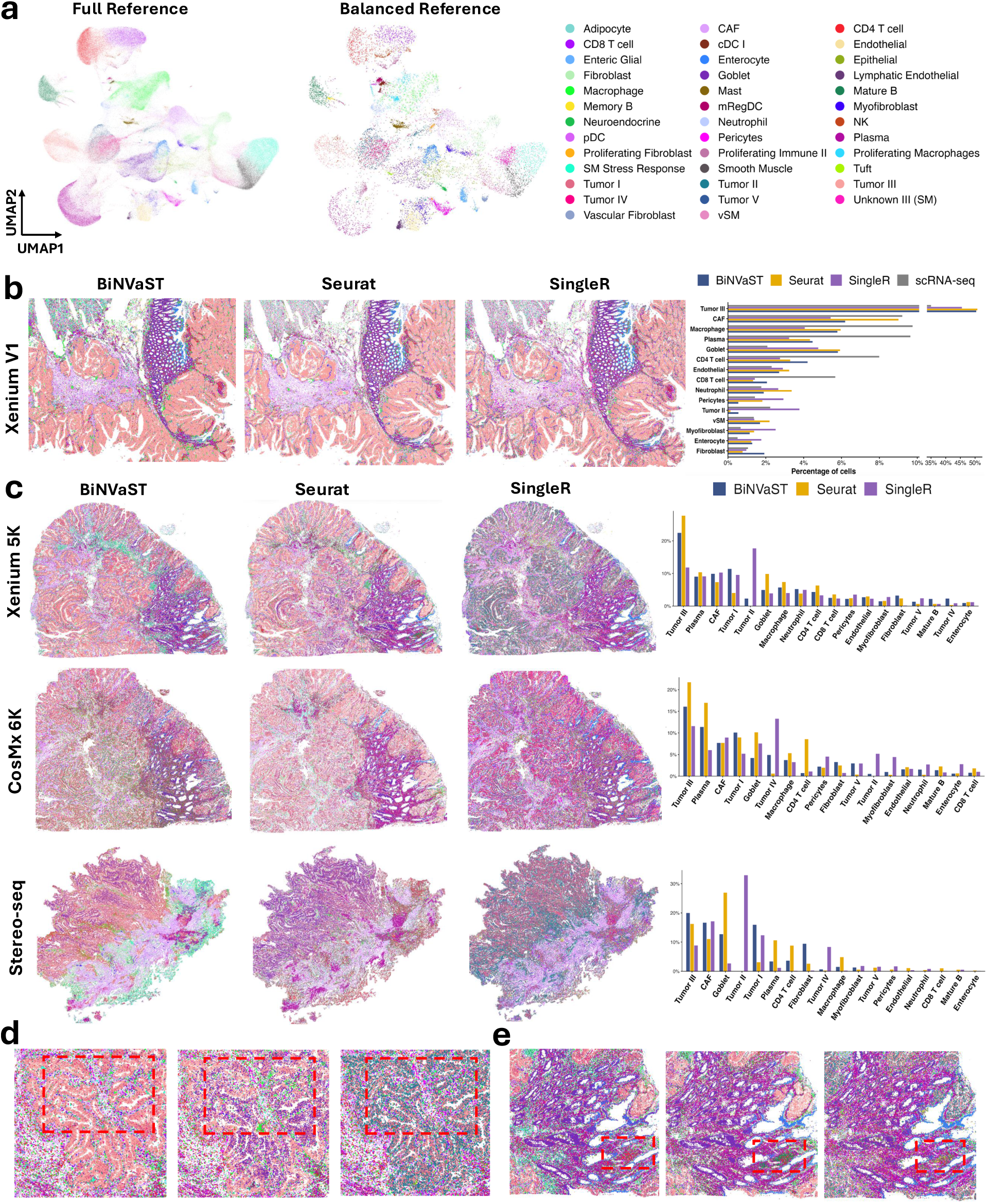
Predicted cell labels across spatial platforms in CRC. **(a)** UMAPs of the full and cell-type-balanced CRC scRNA-seq references, with pie charts summarizing their cell-type compositions. Colors indicate cell types and are shared across panels. **(b)** Spatial cell-type maps of the CRC Xenium V1 sample produced by BiNVaST, Seurat, and SingleR, and cell proportions of leading cell types for each method together with the tissue-matched scRNA-seq reference. **(c)** Spatial annotations and cell-type proportions for the CRC Xenium 5K, CosMx 6K, and Stereo-seq samples. For each platform, maps are shown for BiNVaST, Seurat, and SingleR using color palette in panel (a). **(d-e)** Enlarged representative tissue regions comparing the local cell-type organization recovered by BiNVaST (left), Seurat (middle), and SingleR (right). Red dashed boxes mark regions of interest used for visual comparison.

We compared BiNVaST with Seurat, SingleR, and CellTypist on multiple sections from another CRC tumor tissue [15], profiled by Xenium 5K, CosMx 6K (FFPE) and Stereo-seq (fresh frozen), respectively. The above CRC scRNA-seq data with *n* = 226, 856 cells was used as external reference to compare four methods. Machine learning algorithms like BiNVaST and CellTypist usually requires a large number of reference (or synthetic) cells for model training, while the performance of Seurat (SVD-based) and SingleR (correlation-based) algorithms may vary by the cell proportions in reference. Hence, we also evaluated Seurat and SingleR using the subset of *n* = 16, 564 cells, with results summarized in Supplementary File and visualized in Supplementary Figures S2-S7.

Overall, the performance of each methods on was better in Xenium and CosMx compared to Stereo-seq (Supplementary Figure S8-S25). With the full reference, Seurat showed concordance across Xenium and CosMx in annotating the major regions (Figure 4c) predominated by Plasma, Goblet, Enterocyte cells, Tumor III malignant cells, Cancer-Associated Fibroblast (CAF), while CD4+ and CD8+ T cells, Mature B cells, and Tumor I malignant cells exhibited partial agreement between two platforms (Supplementary Figures S14, S17, S15, S18). Interestingly, both BiNVaST and Seurat identified more Tumor I malignant cells (11.23%, 8.96%) in CosMx 6K than Xenium 5K (7.93%, 3.99%), and detected Neutrophil-enriched domains only in Xenium. Plasma cells were concordantly annotated by BiNVaST across Xenium and CosMx, while Seurat overestimated plasma cells (16.95%) in CosMx compared to Xenium. On the other hand, BiNVaST consistently identified divergent tumor cell states (i.e., Tumor I & III) on both platforms.

Rare malignant cell states Tumor IV & V were identified by BiNVaST across two platforms, which were barely detected by Seurat. The regions enriched with other normal cell populations were robustly annotated as well (Supplementary Figures S8, S9), although the abundance of immune cells was slightly under-estimated compared to Seurat (Supplementary Figures S14, S15). Moreover, in the Stereo-seq section, all three methods annotated the regions enriched by CAF, but Seurat and SingleR overestimated Goblet cells (Supplementary Figures S10, S16, S22). CellTypist was applied to each section with the raw vs. majority-voted submodules. Each section was poorly annotated by this tool ((Supplementary Figure S26), especially in CosMx, although Xenium 5K annotation was partially correct (e.g., CAF, Plasma, Goblet) compared to the results from competing methods. The CellTypist majority-voted output assigned 76.9% of CosMx cells to Myofibroblast and 16.8% to Tumor II, leaving little spatial or biological diversity. Stereo-seq also showed collapse in CellTypist majority-voted annotation, with 42.7% Tumor I and 30.5% CAF, although the predicted labels detected the niches or domains not captured by raw labels (Supplementary Figure S26). BiNVaST consistently predicted Tumor III (*>*10%) and Tumor I (*>*5%) across three platforms, whereas Tumor III was the leading malignant cell state. Goblet was reasonably annotated by BiNVaST in Xenium and CosMx (*<* 5%), but its abundance was moderately inflated in Stereo-seq (12%). Seurat overestimated Goblet in Xenium (10%) and Stereo-seq (26%). The performance of BiNVaST in CosMx and Stereo-seq was less ideal compared to Xenium, whereas Tumor IV was moderately overestimated in CosMx and under-estimated in Stereo-seq. This malignant cell state was barely recovered by Seurat across three platforms. Seurat stably annotated CD4 T cells across platforms, while BiNVaST and SingleR missed this immune cell type in either CosMx or Stereo-seq. Further, we visualized the niches defined by CAF, Myofibroblast, Macrophages, Goblet, Enterocyte within two ROIs in Figure 4d-e. Obviously, in the first ROI, the boundaries of CAF-enriched niches predicted by BiNVaST are better defined with primary malignant cells (Tumor III & I) well annotated, outperforming Seurat and SingleR. In the second ROI, all three methods robustly captured the regional distributions of Goblet versus Enterocyte in colon tissue. Seurat identified a niche dominated by Mature B cells with randomly co-localized CD4 T cells, while BiNVaST labeled this domain with more CD4 T cells and scatter Mature B cells.

### Quantitative concordance evaluation in CRC sections across platforms

Cross-platform consistency was further assessed by correlating cell-type-specific gene reference-target concordance scores, i.e., Rank-Biased Overlap (RBO), Jaccard Index, and Weighted Jaccard Index, between each pair of spatial platforms. After abundance filtering, 18, 11, and 12 shared cell populations were retained for the Xenium–CosMx, Xenium–Stereo-seq, and CosMx–Stereo-seq comparisons, respectively. Definitions for these concordance scores are described in Methods. For each platform pair, BiNVaST achieved higher or comparable mean Spearman correlations in Weighted Jaccard than Seurat and SingleR (Figure 5a), with the highest mean cross-platform consistency by Jaccard and relatively low variation across platform pairs, while performance remained comparable to the benchmark methods across the three metrics.

**Fig. 5.**
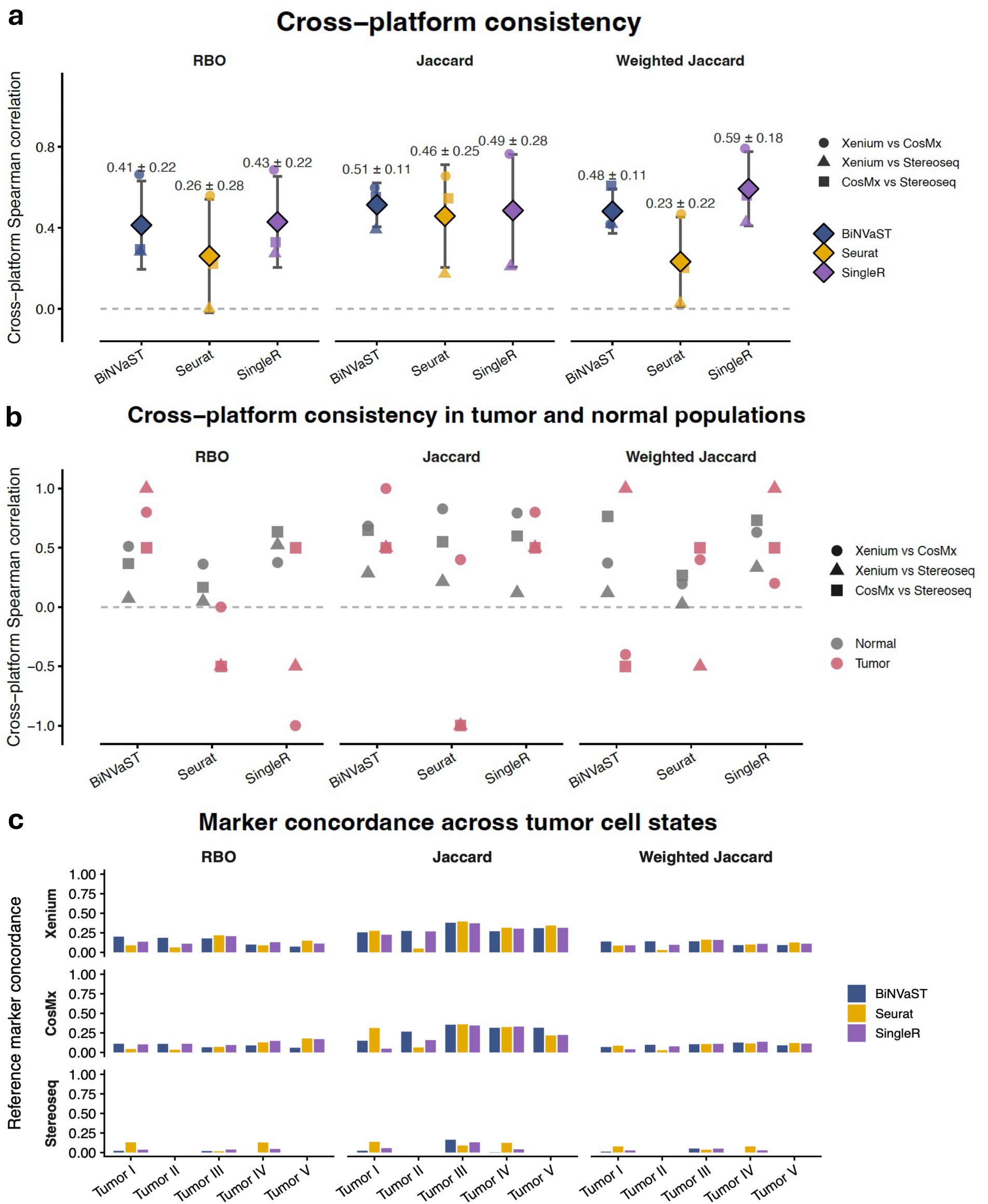
Cross-platform concordance in cell-type-specific genes. **(a)** Cross-platform consistency of BiNVaST, Seurat, and SingleR annotations assessed using rank-biased overlap (RBO), Jaccard similarity, and weighted Jaccard similarity. Spearman correlations are shown for each platform pair (Xenium-CosMx, Xenium-Stereo-seq, and CosMx-Stereo-seq); diamonds and error bars summarize the mean and variability across platform comparisons. **(b)** Cross-platform consistency stratified into tumor and normal cell populations. **(c)** Concordance between spatially derived and reference marker rankings for four tumor cell states (Tumor I-IV) in Xenium, CosMx, and Stereo-seq. Bars show RBO, Jaccard, and weighted Jaccard concordance for BiNVaST, Seurat, and SingleR.

We next stratified the cross-platform analysis into tumor and normal populations (Figure 5b). Tumor-specific correlations were more variable than correlations based on normal populations, consistent with the smaller number of eligible tumor states available for correlation analysis (3–4 tumor states versus 8–14 normal populations per platform pair). For BiNVaST, tumor-state correlations remained positive across all three platform pairs for both RBO (0.50–1.00) and Jaccard (0.50–1.00), whereas weighted Jaccard correlations were more variable. Overall, the relative tumor-versus-normal pattern depends on the platform pair and metric, with no uniform shift observed across annotation methods.

Finally, we examined reference concordance for individual tumor states (Figure 5c). Tumor I, III, and IV met the abundance threshold and were evaluated, whereas Tumor II was excluded for Seurat and Stereo-seq because this state was not successfully identified. Across the retained tumor states, BiNVaST showed concordance with reference within the range of Seurat and SingleR, with several state-specific strengths. For example, BiNVaST showed the highest RBO for Xenium Tumor I (0.201, compared with 0.090 for Seurat and 0.136 for SingleR) and the highest Jaccard for Stereo-seq Tumor III (0.164, compared with 0.090 and 0.131, respectively). No single method consistently achieved the highest concordance across all tumor states, platforms, and metrics.

### Pediatric High-Grade Glioma: Xenium 5K

We applied BiNVaST, Seurat, and SingleR to a newly diagnosed pediatric High-Grade Glioma (pHGG) tumor using pre-annotated scRNA-seq reference from 19 tumor samples matched by molecular subtype, i.e., H3 G34-mutant Diffuse Hemispheric Glioma (DHG). In this single cell reference, we defined seven malignant cell states, i.e., Astrocyte (AC)-like, Mesenchymal (MES)-like, Radial Glia (RG)-like, Neuronal Progenitor Cell (NPC)-like, Interneuronal Precursor Cell (INPC)-like, early GABAergic Interneuron (eIN)-like, Oligodendrocyte Precursor Cell (OPC)-like, and six non-malignant (sub)populations, i.e., microglia, M1 macrophage (pro-inflammatory), M2 macrophage (immunosuppressive), monocytes, mature oligoden-drocytes, endothelial cells. The predicted cell labels are visualized by spatial map (Figure 6a), while the molecular embeddings predicted by the optimized MVAE branch in BiNVaST are visualized by principal components (PCs) and UMAPs, illustrating the molecular difference among cell populations. The synthetic molecular embeddings from the training step visualized by PCs and UMAPs (Figure 6b) illustrated the robustness of built-in simulator that mimics zero-inflation in query Xenium profile and the validity of Optimal Transport that pairs spatial coordinates and synthetic cells.

**Fig. 6.**
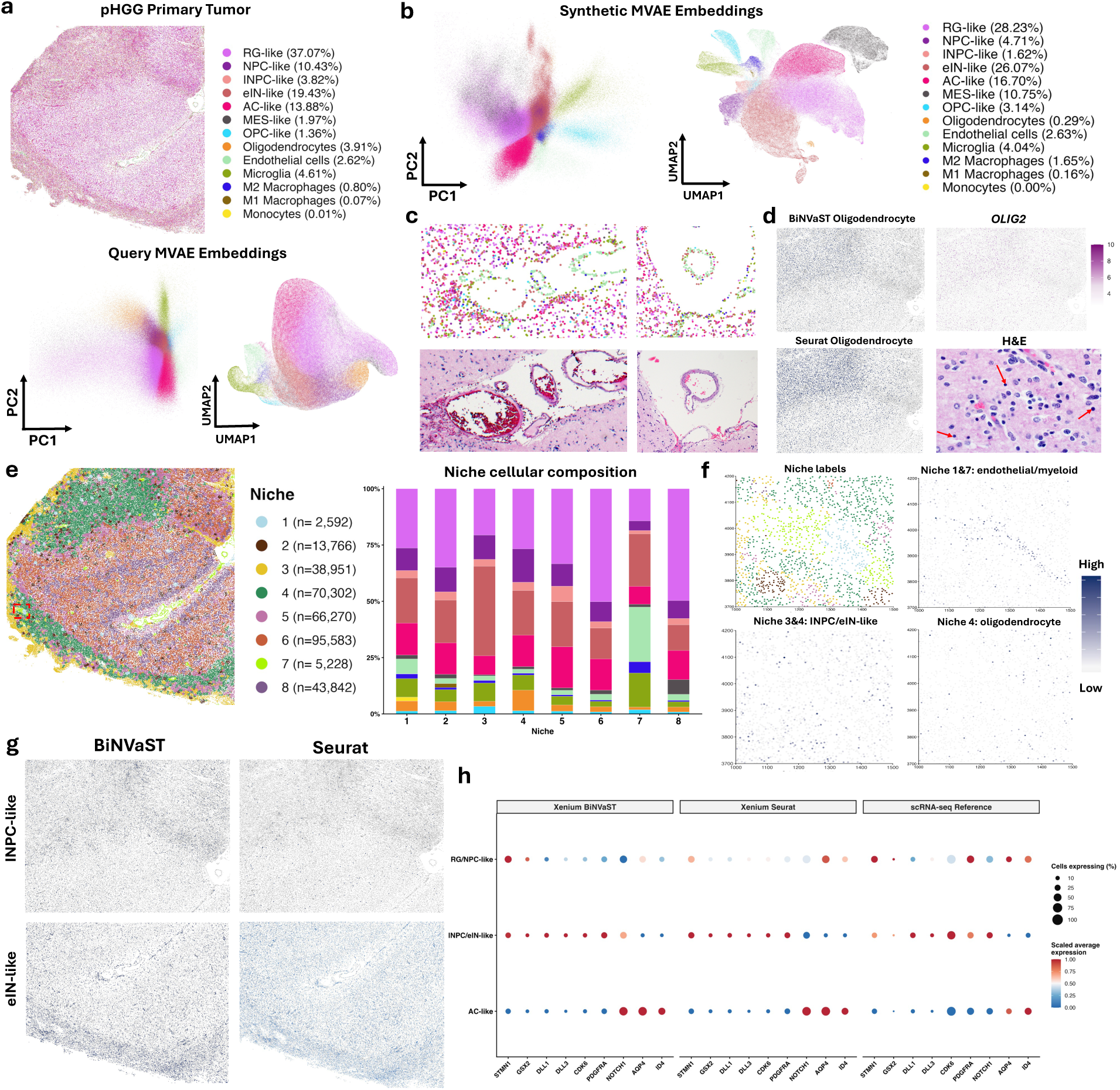
BiNVaST resolves tumor microenvironment and spatial niches in pediatric high-grade glioma. **(a)** BiNVaST cell-state map of the primary pHGG tissue and PCA/UMAP representations of the target molecular embeddings. Cell-type proportions are reported in the legend. **(b)** PCA and UMAP representations of the synthetic molecular embeddings used for model training, colored by cell identity. **(c)** Representative ROIs showing the spatial arrangement of predicted cell states alongside the corresponding histology. **(d)** Spatial localization of oligodendrocytes predicted by BiNVaST and Seurat, *OLIG2* expression, and the corresponding H&E region; arrows mark cells with oligodendrocyte-like morphology. **(e)** Eight spatial niches identified from local cellular neighborhoods and their mean cell-type compositions. The stacked bars show the proportional contribution of each cell type to each niche. **(f)** Cell-level niche labels in an ROI and spatial feature maps of weighted marker scores for endothelial/myeloid-enriched Niches 1 & 7, INPC/eIN-like-enriched Niches 3 & 4, and oligodendrocyte-enriched Niche 4. Darker color indicates a higher weighted marker score. **(g)** Spatial maps comparing BiNVaST and Seurat assignments for INPC-like and eIN-like cells. **(h)** Bubble plot comparing selected neural-development and lineage markers across BiNVaST, Seurat, and the scRNA-seq reference. Rows represent the combined RG/NPC-like, INPC/eIN-like, and AC-like states.

Overall, BiNVaST (Figure 6c) and Seurat (Supplementary Figure S27) annotations aligned with major anatomic regions (i.e., blood vessels, white matter, and cerebral cortex), while SingleR (Supplementary Figure S27) did not identify these regions and failed to distinguish malignant cell states and myeloid subpopulations. Specifically, BiNVaST identified mature normal oligodendrocytes at reasonable abundance with regional clusters (Figure 6d), relative to the marker *OLIG2* expression and oligodendrocytes distribution/distance in H&E. Neoplastic cells in this H3 G34-mutant DHG tumor are OLIG2-negative. Therefore, *OLIG2* expression in this tumor marks non-neoplastic oligodendroglial cells more cleanly than it would in other glioma tumors.

Next, we built niches on cell labels predicted by BiNVaST and Seurat, individually, clustering the cellular composition of each spot’s neighborhood defined by the nearest *K* = 60 spots/cells. The niches identified from BiNVaST labels (Figure 6e) successfully captured domains enriched by endothelial/myeloid cells (Niche 1 & 7), M1 macrophage (Niche 2), oligodendrocyte/NPC-like (Niche 4), eIN-like (Niche 3), INPC-like (Niche 5), and RG-like (Niche 6 & 8), while niches derived from Seurat cell labels did not distinguish malignant cells vs. endothelial/myeloid cells (Supplementary Figure xx). Notably, the niches track along specific anatomic structures in the cerebral cortex and underlying white matter, suggesting their influence on tumor cell state composition. Niche 3 is deep white matter and Niche 4 is subcortical white matter, while Niches 5 & 6 are infiltrated cerebral cortex. Niche 7 is pia/leptomeningeal and Niche 8 is subpial/layers 1-2.

For an ROI dominated by Niches 1, 3, 4, 7 (Figure 6f), we calculated the weighted expression scores of marker genes for endothelial/myeloid in Niches 1 & 7 (*CLDNA, ENG, PECAM1, P2RY12, TMEM119, CD163, CD68, ADGRA2*, INPC/eIN-like tumor cells in Niches 3&4 (*DLL1, DLL3, ASCL1, GAD1*), and mature oligodendrocytes in Niche 4 (*OLIG2, SOX10*), visualized in Figure 6f. For each marker score, we confirmed localized expression (p= 0.0001, 0.0006, 0.0014) and cell-cell heterogeneity (p= 0.0001, 0.0001, 0.0046) by one-sided permutation test on the mean and variance. Localized spatial dependence, quantified by Moran’s I autocorrelation [20] for the weighted marker score, was only observed in endothelial/myeloid niches (p= 0.0001), confirming the frequent endothelial-myeloid colocalization. INPC/eIN-like malignant cells in this ROI did not show significant colocalization, being consistent with the exclusion of Niche 5 in this region. Further, the INPC-like and eIN-like cells predicted by BiNVaST showed distinct regional patterns (Figure 6g), while Seurat yielded either spatially diffuse prediction of INPC-like cells or over-abundant eIN-like cells. The cell state markers or disease-specific driver genes in RG/NPC-like, INPC/eIN-like, AC-like neoplastic cells identified by previous studies [21, 22] were visualized (Figure 6h) for Xenium and scRNA-seq reference. PDGFRA expression in the reference cohort, where at least *n* = 12 tumors (patients) had PDGFRA alteration, was higher in progenitor cell states (RG/NPC-like), while the target sample profiled by Xenium 5K was PDGFRA wild-type and showed higher expression in more differentiated cell state (INPC/eIN-like).

## Discussion

We developed BiNVaST, a niche boundary-preserving and synthetic data-guided variational autoencoder framework for post-segmentation cell phenotyping across high-resolution ST platforms. By integrating user-defined single-cell references with query spatial profiles, BiNVaST addresses a central limitation of existing annotation tools: their dependence on generic or imperfectly matched references that may not capture disease-specific cell states. This feature is especially important in cancer, where malignant programs, immune subpopulations, and spatially organized niches often diverge from normal-tissue atlases.

Across the current benchmarking analyses, BiNVaST showed improved performance over commonly used reference-transfer methods by combining molecular similarity with spatial neighborhood information. In colorectal and ovarian cancer datasets, BiNVaST better preserved local niche boundaries and improved concordance of cell-type assignments across platforms and serial tissue sections. In pediatric high-grade glioma, BiNVaST resolved malignant cell states and non-malignant tumor microenvironment populations that were difficult to distinguish using standard label-transfer approaches. These results suggest that high-resolution ST annotation benefits from models that learn not only from cell-intrinsic expression profiles but also from rare tissue architecture and local spatial organization.

A key strength of BiNVaST is its pre-training strategy guided by a probabilistic simulator, reducing the gap between single-cell reference and spatial measurements by generating platform-adjusted in silico data to reflect the sparsity and spatial structure of the target ST profile. This design allows the model to pretrain on labeled synthetic data before adapting to the query tissue, improving robustness when real spatial data with labels are unavailable. The use of user-defined references further makes BiNVaST flexible for rare diseases, pediatric tumors, and other settings where public atlases are incomplete or biologically mismatched.

The choice of PCA basis is an important component of synthetic-data construction. If the reference and query tissues closely match in expression spaces, fitting PCA to the reference and projecting the target, or fitting PCA to the target and projecting the reference, should preserve similar neighborhood geometry and have little effect on the resulting predictions. In practice, this assumption is often too idealistic because single-cell and spatial datasets can differ substantially in capture chemistry, sequencing depth, gene coverage and sparsity. The selected PCA basis can therefore change the cross-dataset distances used to assign synthetic spatial coordinates and, consequently, alter the final target predictions. This sensitivity further motivates generating target-adjusted synthetic training data instead of training the model directly on the single-cell reference. We recommend reference-based PCA as the default when the reference is less sparse than the target. For target spatial profile with higher expression and/or less sparsity than reference, target-based PCA projection can be a preferred option.

Deep neural network (DNN) hyperparameters, particularly the number of training epochs and batch size, likewise affect convergence, prediction performance and computational cost. The internal-validation strategy described in the Methods provides a label-preserving approach for comparing candidate configurations, but we also encourage users to perform a practical grid search over plausible settings for their own tissue and platform. In the above benchmarking studies, the configuration was determined by the ground truth of certain normal cells, e.g., endothelial cells and blood vessels in H&E, or the abundance/rank of tumor cell states. The trade-off was evident in DHG runs with 2000 input genes and 300,000 cells. Training for 100 epochs with a batch size of 8,192 was completed in 23 minutes, while training for 200 epochs with a batch size of 512 required more than 2.5 hours and yielded the optimal cell labels. Thus, doubling the epoch count while using a substantially smaller batch increased the number of optimization steps and extended runtime by approximately 6.7-fold.

Several limitations remain in BiNVaST. First, Visium HD 2×2 *µ*m bins are highly sparse and may represent partial cells, extracellular RNA, or mixed local signals. Although BiNVaST can model the high-resolution molecular data with cell segmentation, the simulator in current version cannot directly capture transcript in 2×2 *µ*m bins and incorporate spatial adjacency. Cell-state prediction in this setting remains challenging and sensitive to cell size, sequencing depth, and the availability of informative marker genes. Second, cross-platform benchmarking is complicated by differences in gene panels, tissue section alignment and preprocessing. Third, the current framework depends on the quality and granularity of the input single-cell reference; poorly annotated or under-sampled reference populations may still limit prediction accuracy.

Future work will focus on extending BiNVaST in several directions. We will develop a simulator and prediction framework for Visium HD 2×2 *µ*m data by more explicitly accounting for partial-cell capture and local transcript diffusion. We will also expand benchmarking across additional tumor types, platforms, and orthogonal validation modalities, including histology, immunohistochemistry, and multiplexed protein imaging. Finally, future versions of BiNVaST will incorporate uncertainty-aware niche calling, automated reference evaluation, and downstream statistical testing of niche-specific marker expression, cell-cell heterogeneity, and spatial dependence. Together, these extensions will further support BiNVaST as a generalizable framework for reliable cell typing and niche inference in heterogeneous tissues.

## Methods

### The BiNVaST algorithm

We developed BiNVaST as a three-stage framework for spatially informed cell-type label transfer. First, the method selects a target-informed shared gene panel and generates a labeled synthetic spatial dataset by aligning single-cell reference profiles with the target expression distribution. Second, a dual-branch variational autoencoder learns expression and spatial representations from the synthetic data, and a supervised classifier maps the joint representation to cell-type probabilities. Third, the trained model is applied to the target spatial transcriptomic dataset using the same gene order, expression transformation, and graph-construction procedure.

### Target-informed feature selection and preprocessing

Let *X^t,^*^raw^ denote the target raw-count matrix and *X^r,^*^raw^ the labeled single-cell reference matrix. We first restricted the target matrix to cells with positive total library size and to genes shared by the target and reference datasets. Variable features were then selected from the shared-gene target matrix with Seurat `FindVariableFeatures`, using `selection.method = ‘vst’` and `nfeatures = 2000`. The VST procedure was applied directly to raw counts, without running `NormalizeData` beforehand. It models the mean-variance relationship across genes and ranks genes by standardized residual variance. Thus, the default input panel contained the 2,000 most variable target genes among genes measurable in both datasets; the same genes and gene order were used for the target and reference matrices.

For downstream synthetic-data generation, the selected-gene matrices were transformed as log(1 + 10^4^*x_ig_/L_i_*), where *x_ig_* is the raw count for gene *g* in cell *i* and *L_i_* is the total count over the full biological-gene library of that cell rather than only the selected genes. Target coordinates and reference cell-type labels were retained by cell identifier and checked for one-to-one alignment with their respective expression matrices.

### Synthetic data generation by PCA alignment and entropy-regularized transport

Let 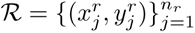 denote the labeled single-cell reference, where 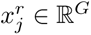 is the expression vector over the selected shared genes and 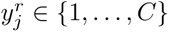 is the annotated cell type. Let 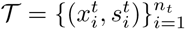 denote the target spatial dataset, where 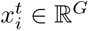 is the target expression profile and 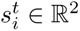 is its spatial coordinate.

We first aligned the reference distribution to the target distribution gene by gene. Positive reference values were rescaled using the corresponding nonzero mode and maximum in the source and target distributions. Zero inflation was then matched with a dropout-only operation. If 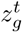 and 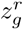 denote the target and reference zero fractions for gene *g*, respectively, a positive aligned reference value was set to zero with probability

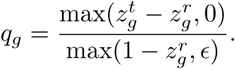

Existing zeros were unchanged. Consequently, additional zeros were introduced only when the target was more sparse for that gene. When the reference was equally or more sparse, *q_g_*= 0 and no nonzero values were backfilled. This one-sided rule prevents synthetic profiles from acquiring expression that was absent from their reference donor cells.

The aligned reference and unchanged target were then embedded in a common principal-component space after centering and scaling both matrices with target-derived gene means and standard deviations. By default, PCA was fitted to the globally aligned reference matrix and the target matrix was projected onto the resulting reference-derived axes. This reference-based PCA is the default because it yields stable axes when the target spatial data are highly sparse. As an alternative for a target dataset with sufficiently dense expression, PCA can be fitted to the unchanged target matrix and the aligned reference can be projected onto the target-derived axes. The two options change only which dataset defines the axes; the same target-derived scaling and subsequent transport procedure are used in both cases.

Let 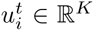 and 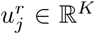 be the target and aligned-reference PCA scores. For each target cell *i* and each cell type *c*, we identified the nearest reference cell within that type,

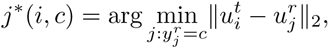

and defined the target-to-type transport cost as

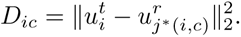

This construction preserves within-type heterogeneity because every cell type is represented by its nearest observed reference cell rather than by a single centroid.

We organized these costs into an entropy-regularized transport plan 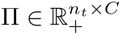. Each target cell supplies one unit of probability mass, and its row of the plan distributes that mass across the *C* candidate cell-type supports. The plan is the solution to

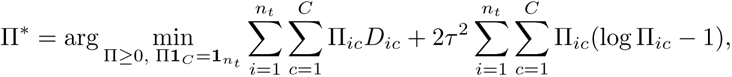

which has the row-wise closed form

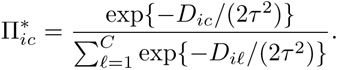

The Gaussian bandwidth *τ* controls the concentration of the plan: smaller values favor the closest type more strongly, whereas larger values preserve greater uncertainty among nearby types. We did not constrain the column marginals of Π. Therefore, the synthetic cell-type composition was allowed to adapt to the geometry of the target dataset instead of being forced to reproduce the cell-type proportions of the reference.

For target cell *i*, the synthetic label was sampled as 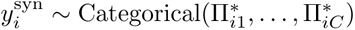. The corresponding nearest reference cell 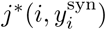 was used as the expression donor. Donor expression was then aligned with cell-type-specific target statistics using the same gene-wise positive-value rescaling and dropout-only zero-rate matching described above. The target coordinate was retained, 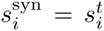, so the synthetic dataset contained exactly one labeled observation for each target coordinate:

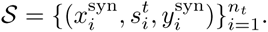

An undirected spatial graph 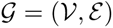 was constructed from the retained coordinates. With the default *k*-nearest-neighbor construction, the adjacency matrix was defined as

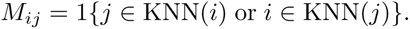

Thus, each synthetic observation combined target-matched expression, an observed target coordinate, local graph context, and a known reference-derived label.

### Dual-branch variational autoencoder and supervised prediction

The model contains expression and graph representation-learning branches. Before training, expression values were transformed with a gene-wise min-max scaler fitted to the synthetic dataset and stored for application to the target data. The expression branch used a multilayer variational autoencoder. For cell *i*, its encoder approximated

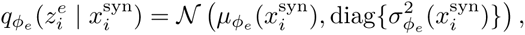

and the decoder reconstructed the scaled expression profile. Its loss combined mean-squared reconstruction error with a Kullback-Leibler penalty toward the standard normal prior. The graph branch used graph-convolutional layers to encode normalized coordinates, local coordinate summaries, and the normalized adjacency matrix. It estimated a graph latent representation 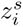 and reconstructed sampled positive and negative edges through inner-product logits 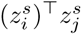. The graph objective combined binary cross-entropy edge reconstruction with a Kullback-Leibler penalty.

The expression and graph representations were concatenated and passed through a nonlinear fused classifier. Expression-only and graph-only classification heads were trained simultaneously as auxiliary objectives. For final prediction, the fused logits were supplemented by a down-weighted graph-head contribution and converted to probabilities with a softmax function. The complete loss therefore combined expression reconstruction, expression and graph Kullback-Leibler terms, graph reconstruction, fused classification, and the two auxiliary classification losses. This design allows the supervised objective learn how expression and spatial information should contribute, while the graph weight controls the residual direct contribution of the graph-only classifier.

### Automatic internal validation and user-directed hyperparameter selection

BiNVaST performs internal validation using only labeled synthetic data; target labels are not used for model fitting or selection. By default, 20% of the synthetic cells are assigned to a contiguous spatial validation crop. A crop center is selected reproducibly from the random seed, and the cells nearest that center form the validation set; all remaining synthetic cells form the training set. A random holdout is also available as an alternative. The spatial crop provides a more stringent assessment than an independent random split because validation cells occupy a held-out tissue region while retaining the same feature-generation mechanism.

During training, validation accuracy and macro-averaged F1 are computed at regular checkpoints. The automatic selection score is

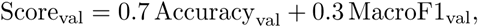

and the checkpoint with the highest score is retained. Macro-F1 ensures that performance on less abundant cell types contributes to selection, whereas accuracy reflects overall classification performance. Training histories, validation metrics, model checkpoints, latent representations, and prediction probabilities are saved for audit and comparison.

The automatic procedure supplies a default model-selection path, but users may also define candidate hyperparameters and select among them. Candidate settings can be evaluated through a grid over quantities such as epoch number, batch size, learning rate, expression and graph latent dimensions, neighborhood size, and loss weights. Continuous training can save exact model snapshots at user-specified epochs, allowing epoch and batch-size combinations to be compared without restarting the same trajectory. Alternatively, users may specify a configuration directly and manually review the saved validation metrics, learning curves, latent-space separation, and spatial prediction maps. The candidate ranges and final settings used in each analysis are reported in the Supplementary Methods.

### Prediction on the target spatial transcriptomic profile

The selected checkpoint and all preprocessing parameters learned from the synthetic data were applied to the target data. The target expression matrix was restricted to the identical ordered gene panel and transformed with the stored expression scaler. A target spatial graph was constructed from its coordinates using the same neighborhood rule. For each target cell, the expression encoder produced 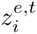 and the graph encoder produced 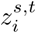. The final target probabilities were calculated by the trained fused and graph classification heads, and the predicted label was

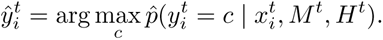

The resulting labels and class probabilities provide a spatially informed transfer of the reference annotations to the target spatial transcriptomic dataset.

### Marker concordance and cross-platform consistency

We quantified concordance between marker genes identified from each annotated spatial dataset and the single-cell reference using three complementary similarity metrics. For each cell type, marker genes from each assay were ranked by decreasing average log2 fold change, and comparisons were restricted to genes shared between assays. A cell type was selected for evaluation if at least 0.1% of cells in a spatial profile were annotated by BiNVaST and a competing method (i.e., Seurat, and SingleR).

Marker concordance was quantified using rank-biased overlap (RBO), Jaccard index, and weighted Jaccard index. RBO compared ranked marker lists while assigning greater importance to agreement near the top of the ranking. The Jaccard index measured unweighted marker-set overlap as the size of the intersection divided by the size of the union. The weighted Jaccard index additionally incorporated marker rank by assigning each gene a weight of 1/r, where r denotes its position in the ranked marker list; similarity was calculated as the sum of the minimum weights across the two lists divided by the sum of their maximum weights. These metrics therefore capture complementary aspects of marker concordance: ranked-list agreement, overall marker-set overlap, and rank-weighted marker overlap, respectively.

To evaluate cross-platform consistency, reference-target concordance scores were first calculated independently for each spatial platform. There are 18 shared populations for Xenium–CosMx, 11 for Xenium–Stereo-seq, and 12 for CosMx–Stereo-seq. For each annotation method and concordance metric, Spearman correlation was calculated between the cell-type-specific scores obtained from the two platforms. In the tumor-versus-normal analysis, the same criteria were used for tumor states and normal populations, individually. The Xenium–CosMx comparison contained 4 tumor cell states and 14 normal cell populations, whereas Xenium–Stereo-seq and CosMx–Stereo-seq contained 3 tumor states and 8 and 9 normal populations, respectively.

### Niche analysis in pediatric high grade glioma

For the DHG Xenium 5K downstream analysis, spatial niches were identified using Seurat’s BuildNicheAs-say function [14]. For each cell, the algorithm defined a local neighborhood using its K spatially nearest cells (calculated by physical coordinates) and represented that neighborhood by the counts of cells per annotated cell type. Cells with similar neighborhood compositions were then grouped into spatial niches by applying K-means clustering, with neighbors.k controlling neighborhood size and niches.k specifying the number of niches. These parameters were determined by the mean composition of each niche (or cluster) and anatomic regions defined by niches. With an optimized configuration, we expect the malignant cell states (e.g., RG-like, eIN-like) and H&E-validated normal cells (e.g., endothelial, oligodendrocytes) are clearly stratified and represented by different niches. The resulting niche assignments and mean cellular compositions were used to identify regions enriched for endothelial/myeloid, oligodendrocyte, neural progenitor-like programs, and rare myeloid subpopulations.

### Single cell reference preprocessing

#### Human normal kidney

The kidney reference was the healthy living-donor kidney single-cell RNA-seq atlas of McEvoy, Murphy *et al.* [23], downloaded as the processed expression matrix and cell metadata from the UCSC Cell Browser (https://cells.ucsc.edu/?ds=living-donor-kidney); raw data are available under GEO accession GSE202109. The downloaded matrix contained 22,484 genes and 27,677 cells from 19 sample libraries. We retained the kidney annotation, `Cell_Types_Broad`, comprising 19 epithelial, stromal, endothelial and immune cell classes. For the shared kidney label vocabulary, PC, CNT and CCD-like were grouped as CCD-like, whereas U1 and U2 were designated Unassigned.

#### Ovarian cancer

The ovarian reference was the matched Chromium Fixed RNA Profiling (Flex) dataset released by 10x Genomics for the Xenium Prime 5K human ovarian cancer specimen [24]. The specimen was a stage IIIB ovarian papillary serous carcinoma, and the Flex material was obtained from FFPE curls adjacent to the sections used for Xenium profiling. The downloaded Cell Ranger v8.0.1 filtered feature–barcode matrix comprised 18,082 measured genes and 17,553 cells. We also downloaded the accompanying barcode-level annotation file, in which the cell identities had been derived from Seurat clustering, marker enrichment and review of the matched Xenium spatial distribution. The HDF5 count matrix was read with `Read10X h5` and used to create an RNA-assay Seurat object with `CreateSeuratObject`. The mitochondrial fraction was calculated and used for QC. Cells were retained when total counts between 200 and 10000, and no more than 10% mitochondria. The resulting reference contained 17,050 cells assigned to 16 cell types.

#### Colorectal cancer

The colorectal cancer (CRC) reference was the 10x Genomics Flex dataset generated for the Visium HD CRC study of Oliveira *et al.* [25]. Processed data are available from the 10x Genomics CRC data portal (https://www.10xgenomics.com/platforms/visium/product-family/dataset-human-crc), raw data are deposited under GEO accession GSE280318, and barcode-level annotations were obtained from the accompanying repository (https://github.com/10XGenomics/HumanColonCancer_VisiumHD/tree/main/MetaData). The aggregated Flex matrix contained 18,082 genes and 279,609 cells from eight CRC tumor and normal-adjacent specimens. The downloaded metadata provided patient and probe-barcode identifiers, a QC flag and hierarchical cell-type labels; the level-2 annotation was used as `cell_type`.

Counts were read with `Read10X_h5` and supplied to CreateSeuratObject We calculated mitochondrial fraction and defined the cutoff as the across-cell mean plus one standard deviation. Cells were retained by total counts and mitochondrial fraction did not exceed this cutoff, leaving 239,837 cells. The final reference contained 226,856 cells assigned to 38 level-2 cell types. The final RNA assay was processed with NormalizeData, FindVariableFeatures, ScaleData in Seurat for visualization. The downloaded level-2 annotations, rather than the de novo cluster numbers, were used by each benchmarked method.

#### Pediatric high-grade glioma

The pediatric high-grade glioma reference was a disease-matched single-cell and single-nucleus RNA-seq atlas comprising 49,392 cells from 19 H3 G34-mutant diffuse hemispheric glioma tumors. The atlas combined a nine-tumor cohort from GEO GSE162989 with ten additional tumors represented in the processed GEO GSE210567/GSE210568 collection [21, 22, 26–28]. Malignant cells were harmonized into RG-like, NPC-like, INPC-like, eIN-like, AC-like, MES-like, OPC-like and cycling programs, while the non-malignant compartment included myeloid, oligodendrocyte, endothelial and astrocyte populations.

Each cohort was processed in Seurat with `NormalizeData`, `FindVariableFeatures` and scaling of all detected genes before PCA. We used 2,000 variable features for the initial malignant-state reference, 5,000 for the ten-sample cohort and malignant-cell reclustering, and 10,000 for the final integrated atlas. Seurat label transfer used PCA dimensions 1-30 for the first cohort and 1-10 for malignant cells in the second cohort; the ten-sample neighbor graph and UMAP used dimensions 1–10 with clustering resolution 2.0, and macrophage/monocyte subclustering used dimensions 1–10 with resolution 1.2.

### Other competing methods

#### Seurat cell label transfer

Cell labels were transferred with Seurat v5’s sketch-based workflow. The reference was again prepared separately for each platform — subset to the panel genes, then re-normalized, re-did HVG and PCA from scratch — and, Each query spatial profile was reduced to a 50,000-cell sketch (12–17% of cells) by Seurat leverage score method. Anchors were identified by FindTransferAnchors with method specified as either PCA or CCA (Canonical Correlation Analysis). Labels were assigned with TransferData function on the sketch and propagated to all cells with ProjectData function.

#### SingleR

SingleR with the aggregated-reference option was applied to OV and CRC sections profiled by four different platforms. Reference and spatial expression matrices were log-normalized, restricted to their shared genes, and supplied to SingleR with aggr.ref = TRUE, which aggregates cells within each reference cell type to reduce the influence of individual-cell variability and unequal cell-type abundance. Spatial cells were classified in computationally manageable batches using the trained reference model. Low-confidence assignments were identified using SingleR’s pruning procedure and designated as unassigned, while the remaining pruned labels were used for downstream spatial visualization and cell-type proportion analyses.

#### CellTypist

Cell-type labels were transferred from full single-cell reference onto three spatial datasets (CosMx 6K, Xenium 5K, Stereo-seq) with CellTypist, a one-vs-rest logistic-regression classifier. Because each platform measures a different gene panel (5,969, 4,827 and 17,471 genes respectively after intersection with the reference), the reference was subset to each panel before normalization and log1p(CPM/10^4^) was recomputed using only panel genes as the library-size denominator, applied identically at training and prediction time; normalizing over the full reference and subsequently subsetting would leave per-cell library sizes inconsistent with the panel. Models were trained with mini-batch stochastic gradient descent (use SGD=True, mini batch=True, batch number=100, batch size=1000, epochs=10, n jobs=1), with mean-centering disabled in the feature scaler (with mean=False) so that the scaling transform preserves sparsity on the largest panel. Classifier performance was first assessed by five-fold stratified cross-validation (stratified on cell type) run independently for each platform panel and for the triple-intersection panel, reporting accuracy, macro-F1, weighted-F1 and per-cell-type F1; final models were then retrained on all 226,856 reference cells. All cells in each spatial dataset were annotated without QC filtering, and raw per-cell calls were smoothed by CellTypist majority voting over a Leiden over-clustering of a 50-PC neighbor graph built on panel genes, with the over-clustering resolution set by CellTypist’s cell-number-scaled heuristic rather than a fixed value. Raw labels, majority-voted labels, maximum class probability and the full 38-class probability matrix were retained for downstream inspection.

#### Azimuth

Pan-human Azimuth is a reference-mapping classifier trained on single-cell profiles from 23 human tissues and organized into a hierarchical cell ontology (https://satijalab.org/azimuth/). We applied the Pan-human model to the cell-segmented human kidney Visium HD dataset. The Space Ranger v4.0.1 outputs were loaded with `Load10X_Spatial(bin.size = c(8, ‘polygons’), image.name = ‘tissue_hires_image.png’)`; the `Spatial.Polygons` assay contained 148,056 segmented cells. This assay was set as the default and processed with `NormalizeData(normalization.method = ‘LogNormalize’, scale.factor = 10000)` before calling `AzimuthAPI::CloudAzimuth(assay = ‘Spatial.Polygons’, model_version = ‘v1’)`. The returned metadata included hierarchical annotations, broad-to-fine labels and `final_level_confidence`. Azimuth labels were mapped to the common kidney vocabulary before concordance analysis.

#### STHD

STHD assigns cell identities directly to individual 2 *×* 2 *µ*m Visium HD bins without requiring prior cell segmentation [12] (https://github.com/yi-zhang/STHD). From the annotated kidney single-cell reference, raw counts were divided by each cell’s library size and then averaged within each cell type to obtain the cell-type-by-gene reference intensity matrix. Mitochondrial and ribosomal genes were excluded, and informative genes were retained when the reference intensity exceeded 0.000125 in at least one cell type and the cell-type-to-overall mean fold change exceeded 2^0.5^. The reference and spatial matrices were restricted to their common genes.

For each 2-*µ*m bin, STHD models gene counts with a Poisson likelihood conditional on the latent cell type and the reference intensity profile. A cross-entropy term encourages neighboring bins to have similar posterior cell-type distributions; the combined objective is *−*LL + *β* CE. Whole-tissue data were divided into patches, fitted patch-wise and recombined. We used the defaults: `n iter = 23`, `step_size = 1`, `beta = 0.1`, `mapcut = 0.8`, `ref_renorm = FALSE`, and background masking enabled with four grid neighbors, `filtermask nrings = 2` and `filtermask threshold = 51` total counts. The maximum-posterior label was assigned when its probability was at least 0.8; otherwise the bin was marked ambiguous, and background-filtered bins were retained as a separate filtered category. Patch-level results were combined to obtain labels for the full kidney section. For comparison with cell-level Azimuth and BiNVaST calls, STHD labels for 2-*µ*m bin centers falling inside each segmentation polygon were aggregated by majority vote.

## Declarations

## Acknowledgment

We thank the patient and family who consented to tissue banking for research. We thank the St. Jude Spatial Omics Core for Xenium 5K profiling of the pediatric high-grade glioma specimen, the St. Jude Biorepository for specimen procurement, histology, and clinical annotation, the Center for Applied Bioinformatics for computational support, and the St. Jude High-Performance Research Computing Facility for computing resources. We also thank 10x Genomics and the investigators who made the public datasets reused in this study available.

## Author contribution

J.Z., J.C., Q.L. conceived this study. J.Z. developed the algorithm and built Python package. Q.L. advised on algorithm and methodology, designed the benchmarking experiments, curated public data, and supervised tool development. J.C., Neuro-Pathologist and Anatomic Pathologist, led the tissue selection, sectioning, QC, and validation in pHGG, as well as reviewed cell labels for each benchmarking experiment. J.Z., L.Li, L.Liu, Q.L. performed the analyses and visualization. J.P., Director of Spatial Omics Core at St. Jude, supervised pHGG Xenium data profiling. B.F., C.C., G.W. discussed methodology, benchmarking results, and disease interpretation. J.Z. and Q.L. led manuscript writing with contributions from L.Li and J.C. All authors reviewed and approved the manuscript.

## Ethics approval and consent to participate

The pediatric high-grade glioma specimen was obtained through the St. Jude Children’s Research Hospital Biorepository under a tissue banking protocol approved by the St. Jude Institutional Review Board with protocol number TBANK Pro00000728, 25-2269, with written informed consent from the patient’s legal guardian and patient assent as applicable. The study was conducted in accordance with the Declaration of Helsinki. All the other datasets were previously published, publicly available, and de-identified, and their reuse required no additional ethical approval.

## Consent for publication

Not applicable. The manuscript contains no identifiable patient information or images.

## Availability of data and materials

The newly-generated Xenium 5K data will be publicly available upon publication in the journal. Public data in this study are listed in Methods.

## Code availability

The source code for BiNVaST is available at https://github.com/zhangjx1996/BiNVaST, accessible upon request. We provide R code for gene selection and input-data preparation, together with Python code for reference-to-query distribution alignment, synthetic-data generation, model training, and final cell-label prediction.

