## Supplementary File for "BiNVaST: niche boundary-preserving and synthetic data-guided variational autoencoders for concordant inference across high-resolution spatial transcriptomic platforms"

Jingxuan Zhang, Lei Li, Limeng Liu, Brooke Fridley, Cheng Cheng, Jasmine
Plummer, Gang Wu, Jason Chiang\*, Qian Li\*

### 1 Ovarian Cancer Xenium 5K sample annotated with internal reference

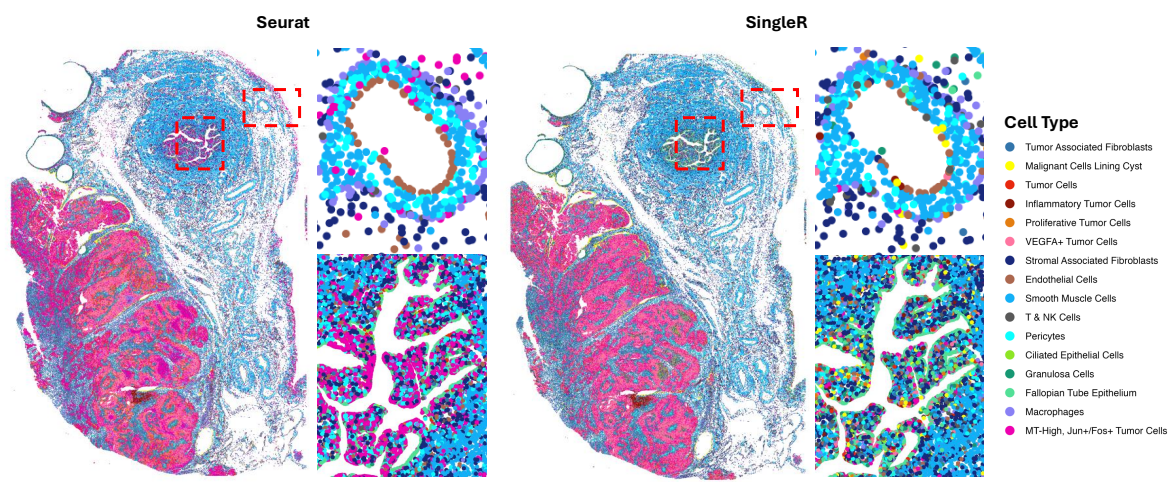

Fig. S1: Spatial maps for cell labels in Xenium 5K ovarian cancer specimen predicted by Seurat and SingleR.

#### 2 Seurat and SingleR with balanced single cell reference

In Xenium 5K, these methods were moderately concordant at the fine-label level. In Seurat results, Tumor III was the most abundant tumor cell states in all three platforms, accounting for 20.1% of Xenium cells, 18.0% of CosMx cells, and 15.2% of Stereo-seq cells. The Seurat annotations were similar between CosMx and Stereo-seq, with a Jensen-Shannon divergence of 0.081, followed by Xenium versus CosMx at 0.088 and Xenium versus Stereo-seq at 0.160. On the other hand, the abundance of tumor cell states (i.e., Tumor I, II, III, IV) predicted by SingleR were severely discordant between three platforms: Xenium (9.5%, 16.2%, 10.4%, 1%), CosMx (4.2%, 9.1%, 9.3%, 12.8%), Stereo-seq (12.9%, 34%, 6.4%, 8.6%). This suggests that CosMx and Stereo-seq shared more similar cell types, while Xenium yields stronger concordance for a subset of epithelial and malignant cells. Details of Seurat and SingleR results using the balanced reference are visualized below.

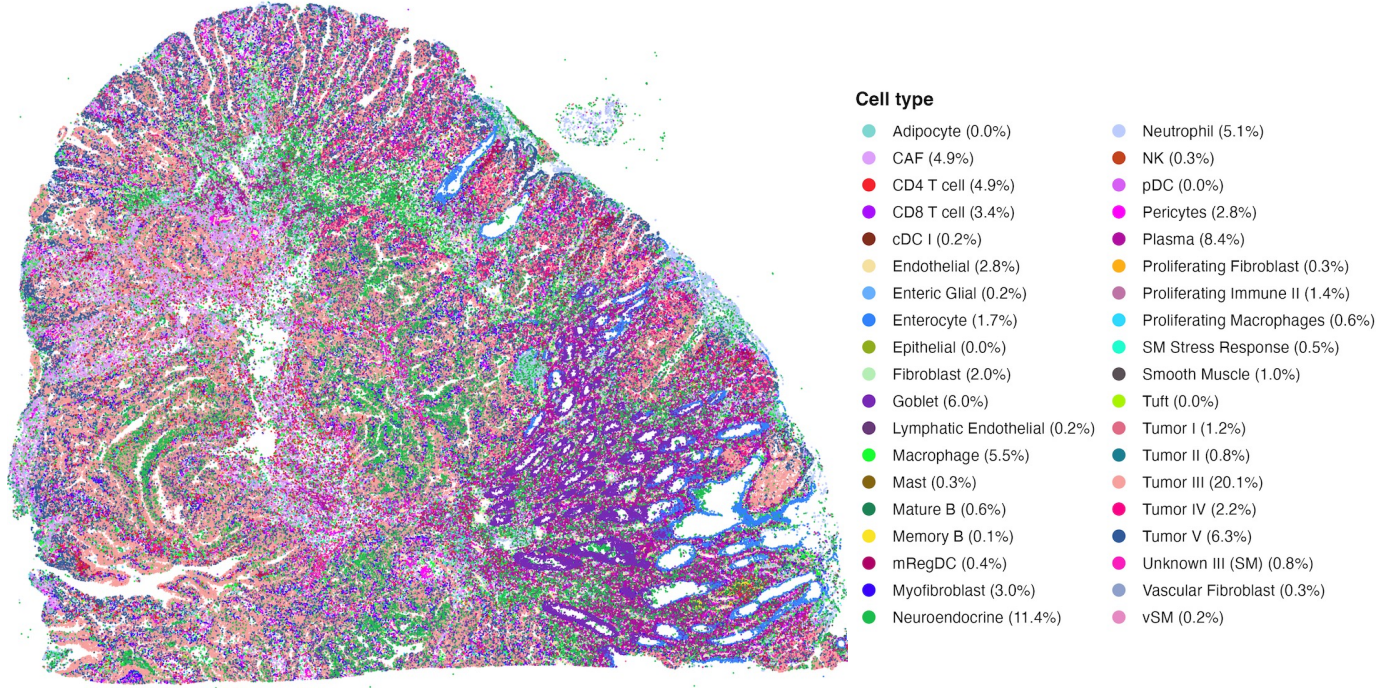

Fig. S2: Spatial map for cell labels by Seurat using balanced reference in Xenium

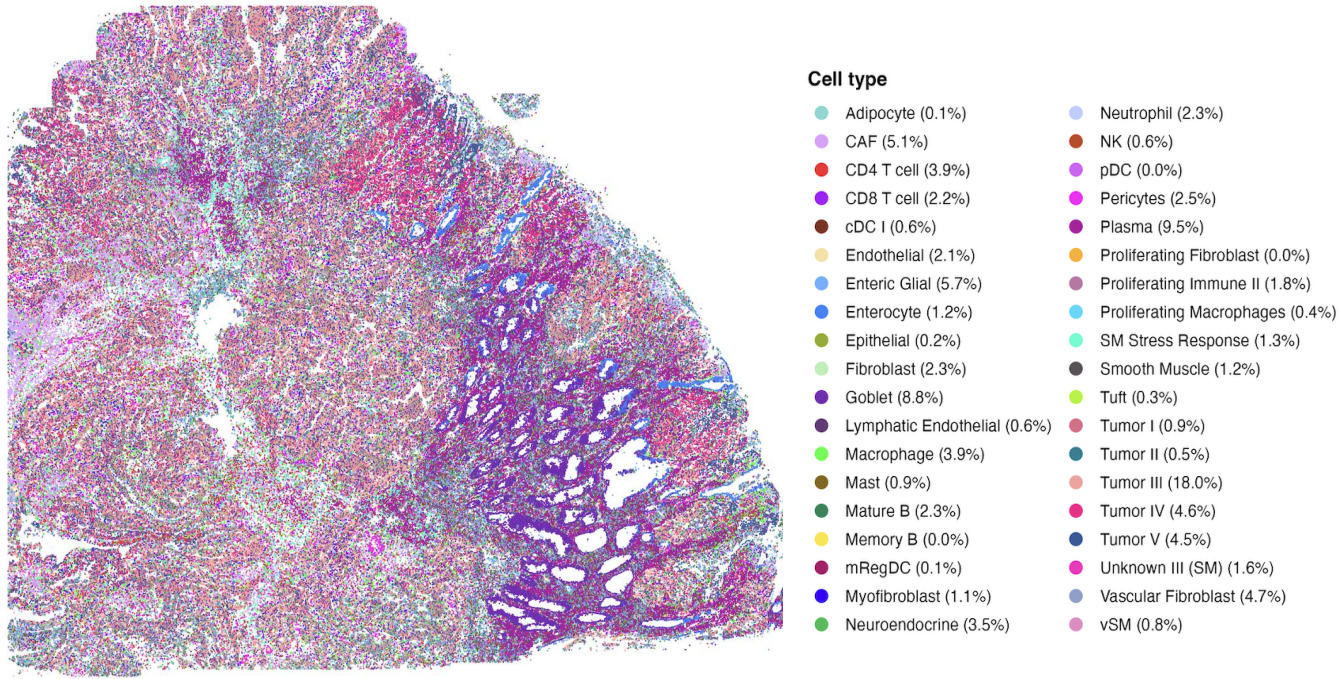

Fig. S3: Spatial map for cell labels by Seurat using balanced reference in CosMx

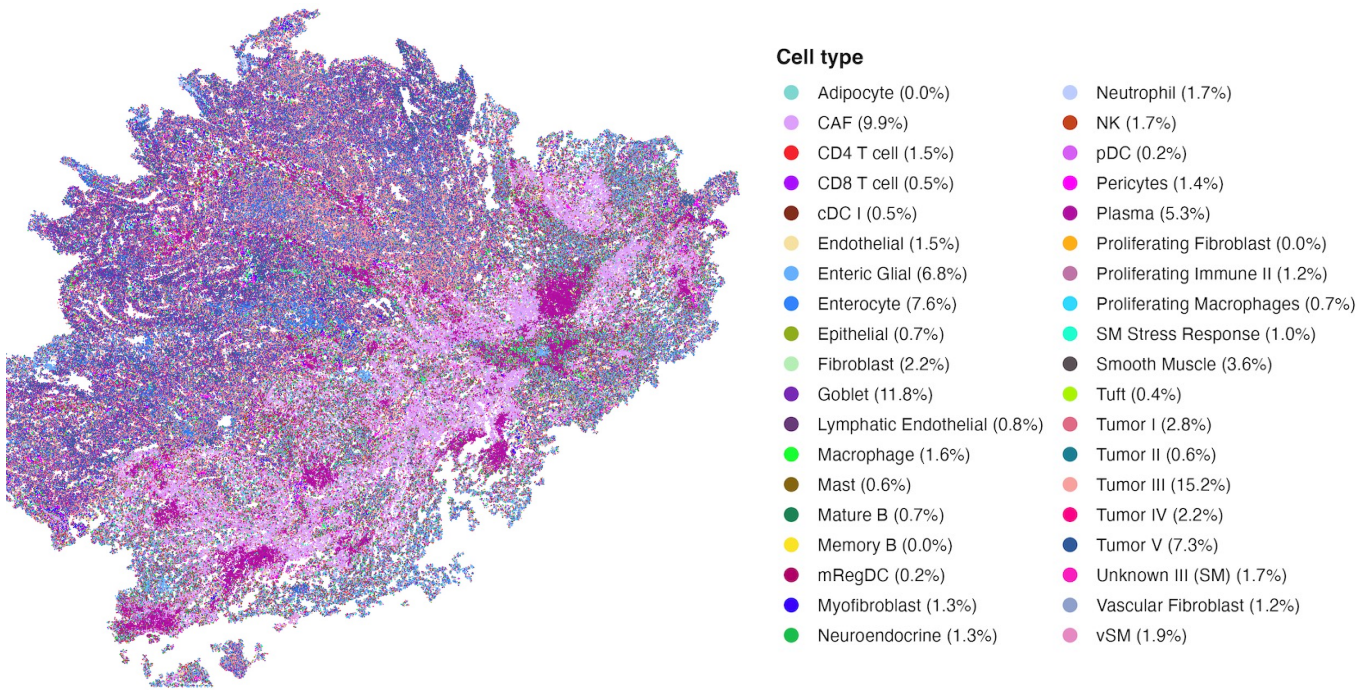

Fig. S4: Spatial map for cell labels by Seurat using balanced reference in Stereo-seq

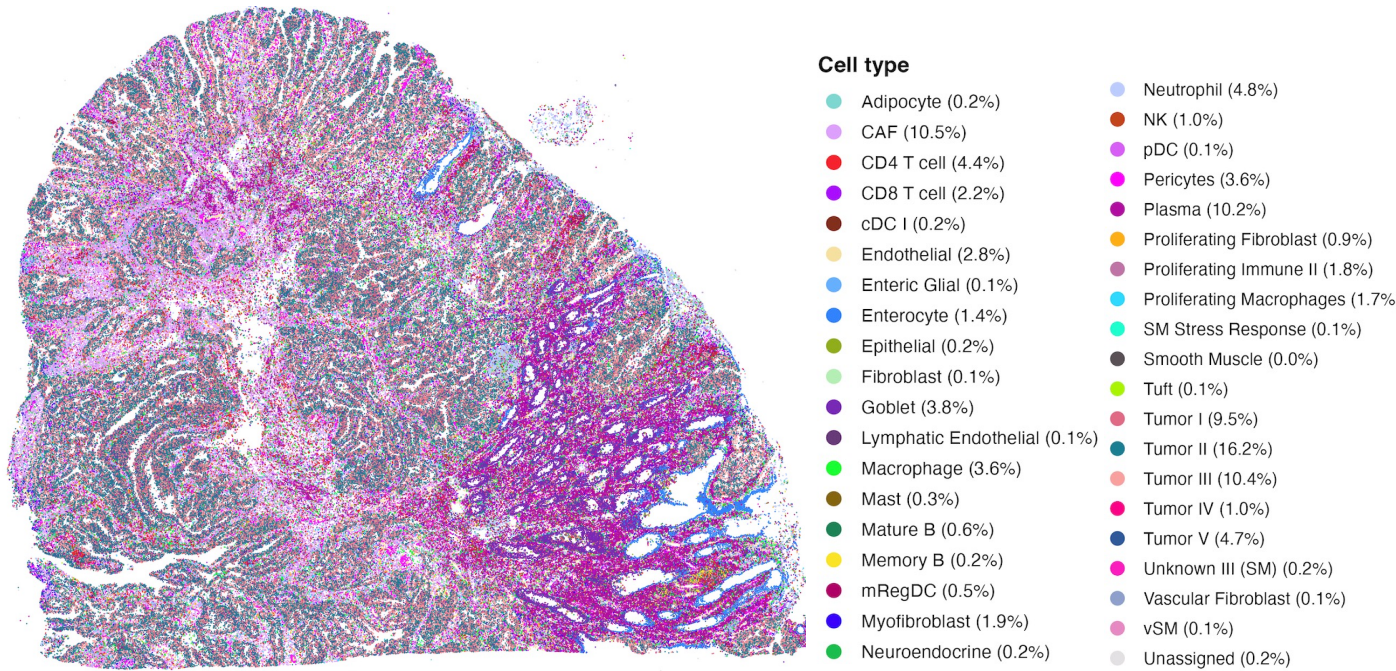

Fig. S5: Spatial map for cell labels by Seurat using balanced reference in Xenium

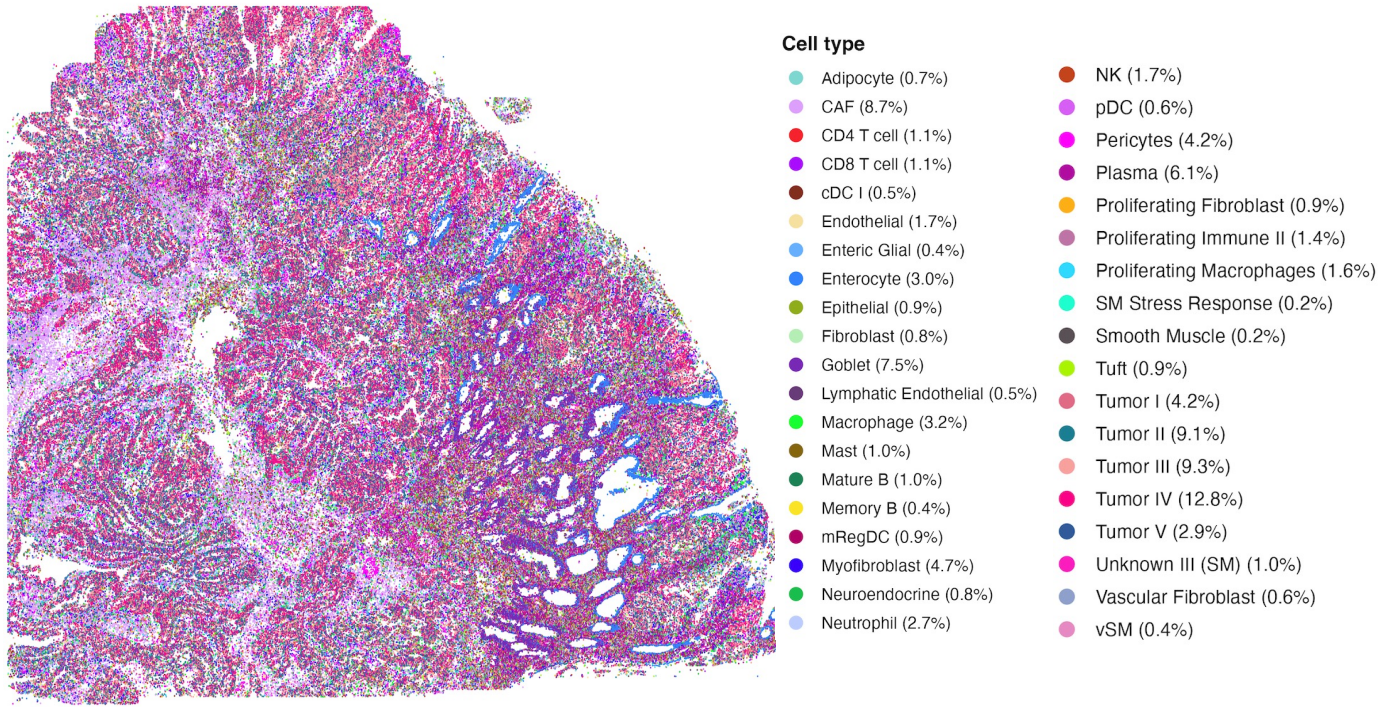

Fig. S6: Spatial map for cell labels by SingleR using balanced reference in CosMx

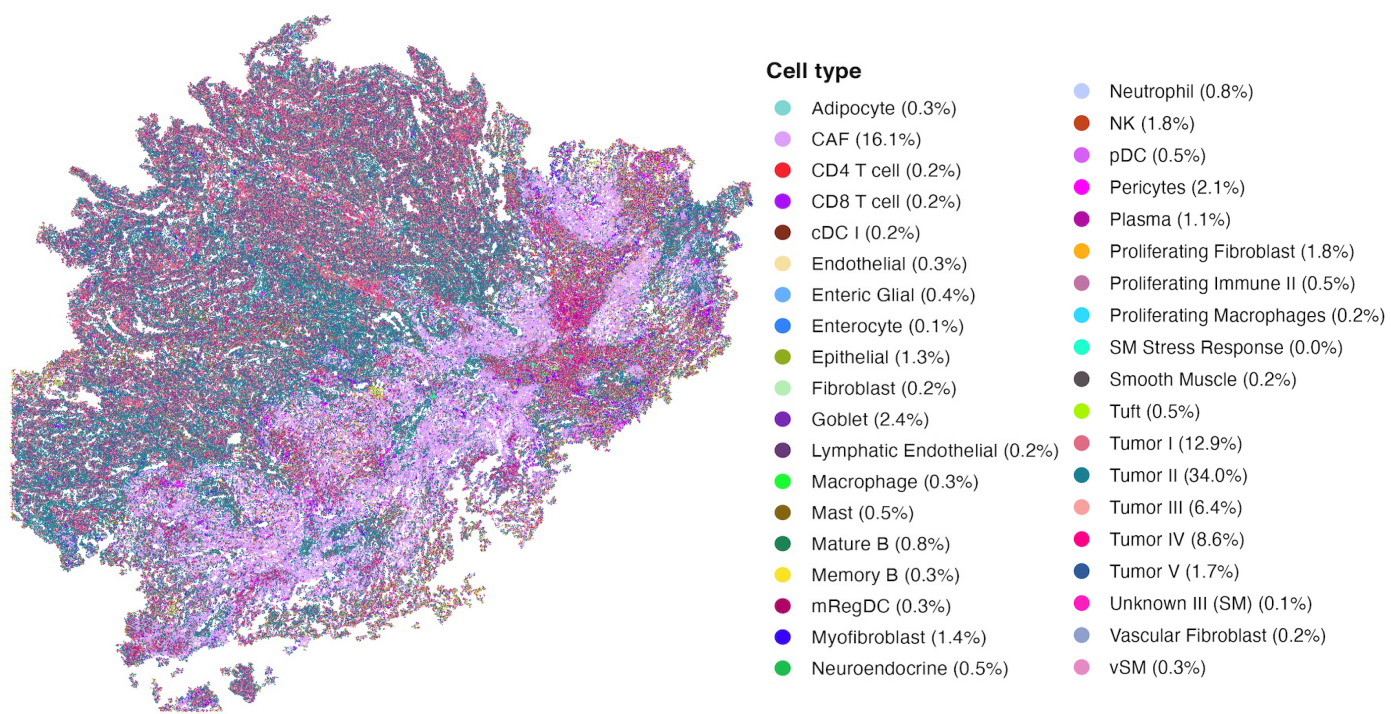

Fig. S7: Spatial map for cell labels by SingleR using balanced reference in Stereo-seq

##### 3 Colon Cancer spatial map by cell type

###### 3.1 Annotation with full single cell reference

###### 3.1.1 BiNVaST

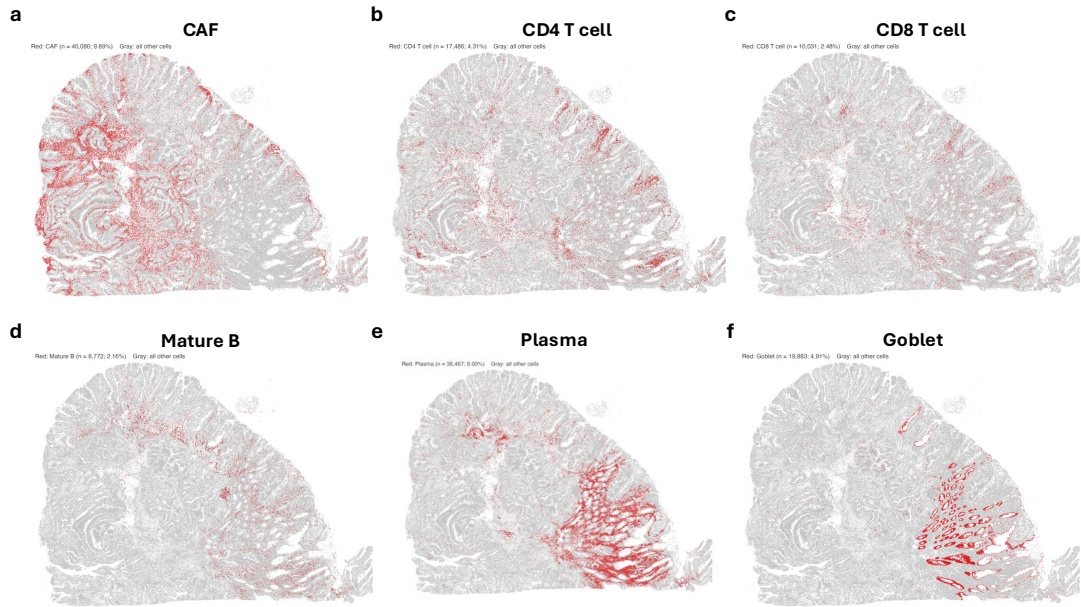

Fig. S8: Individual spatial map for normal cells labeled by BiNVaST in Xenium

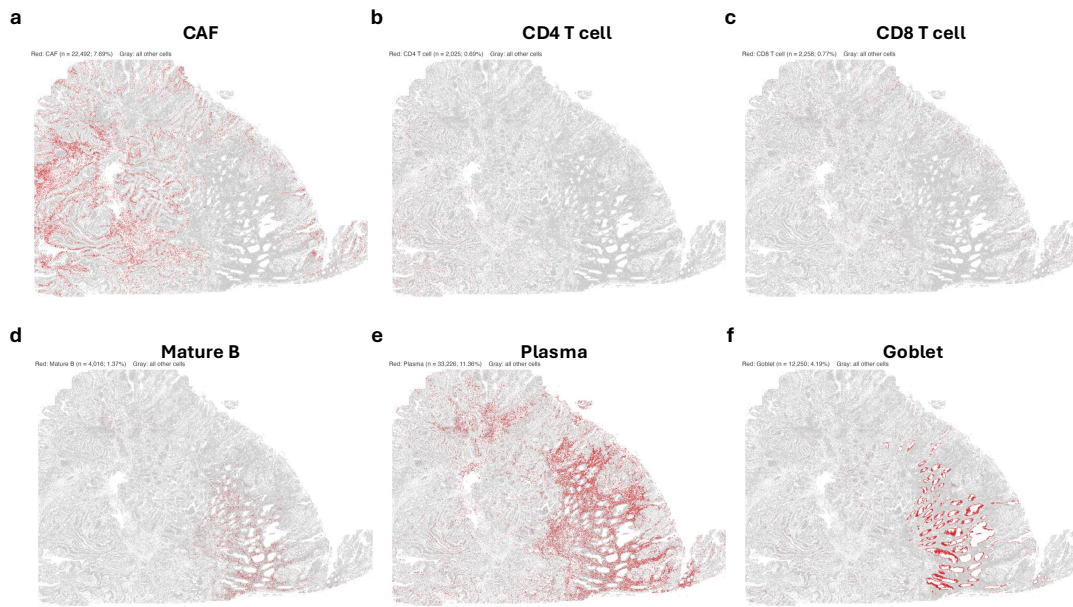

Fig. S9: Individual spatial map for normal cells labeled by BiNVaST in CosMx

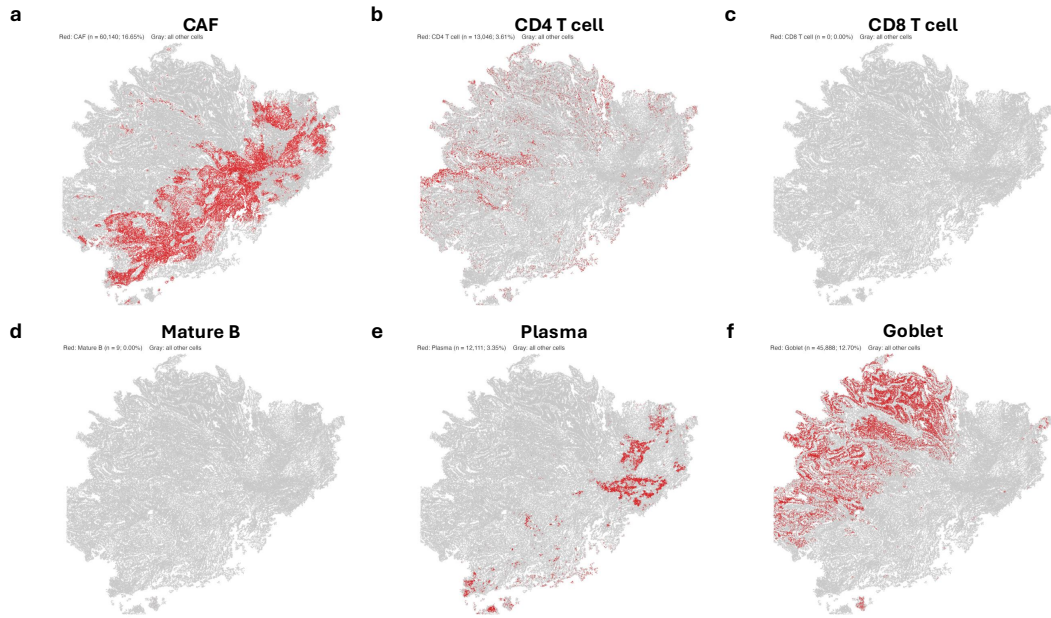

Fig. S10: Individual spatial map for normal cells labeled by BiNVaST in Stereo-seq

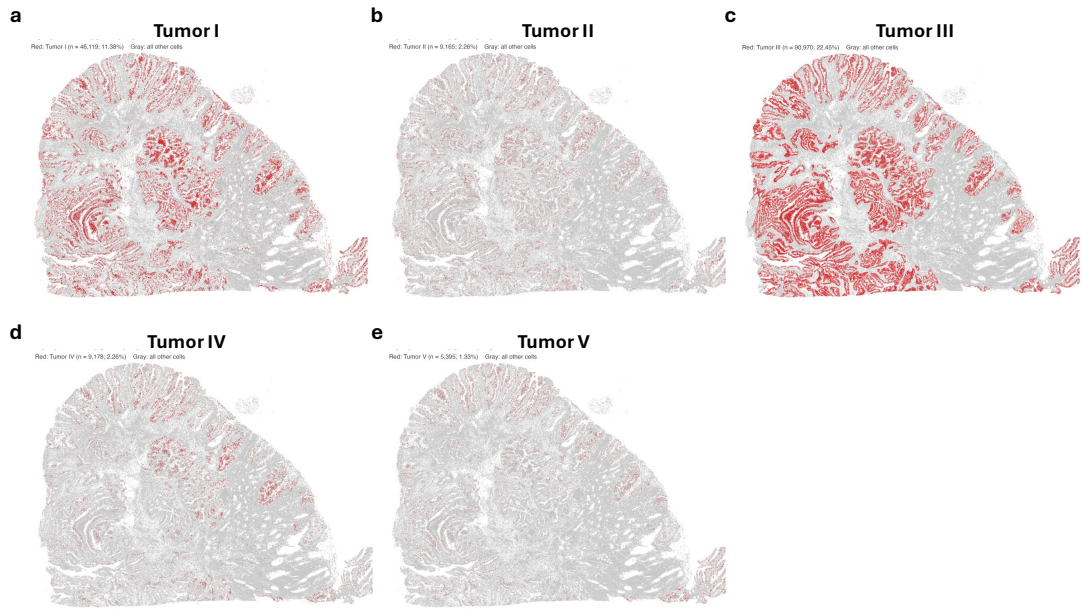

Fig. S11: Individual spatial map for tumor cells labeled by BiNVaST in Xenium

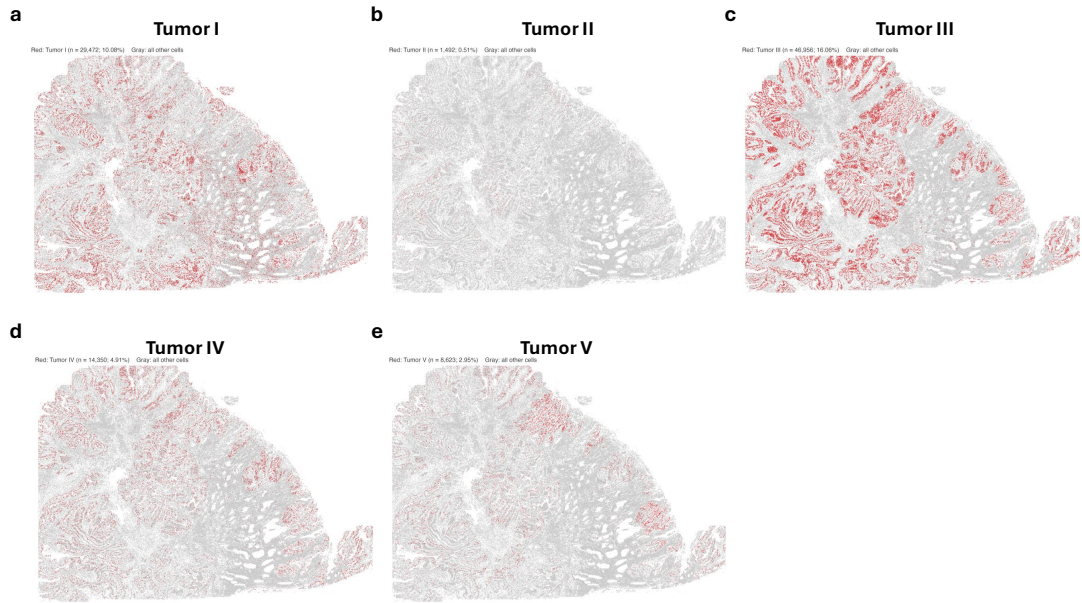

Fig. S12: Individual spatial map for tumor cells labeled by BiNVaST in CosMx

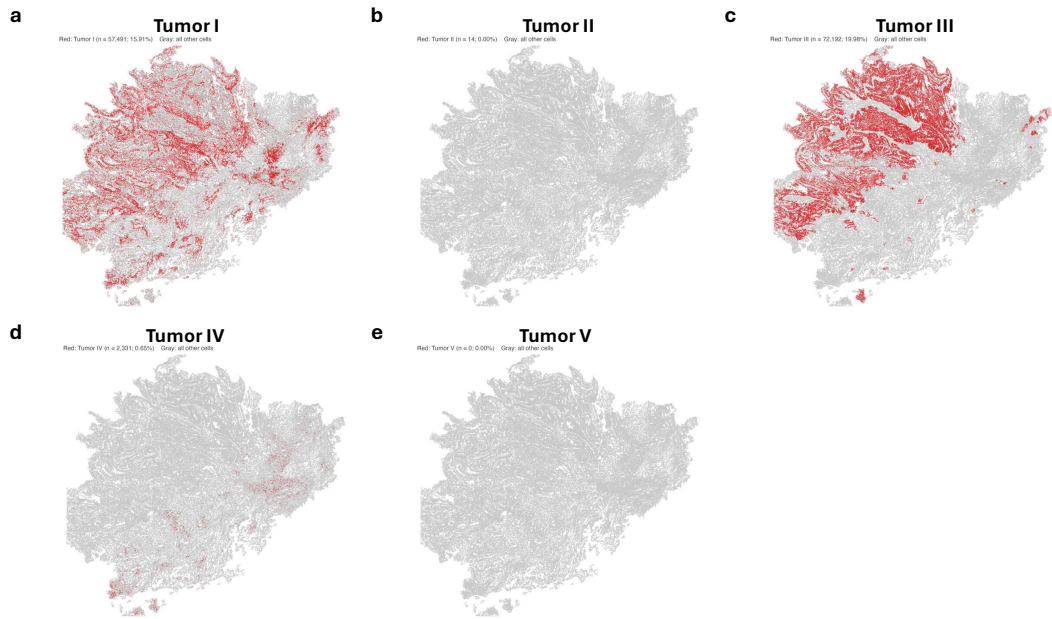

Fig. S13: Individual spatial map for tumor cells labeled by BiNVaST in Stereo-seq

##### 3.1.2 Seurat

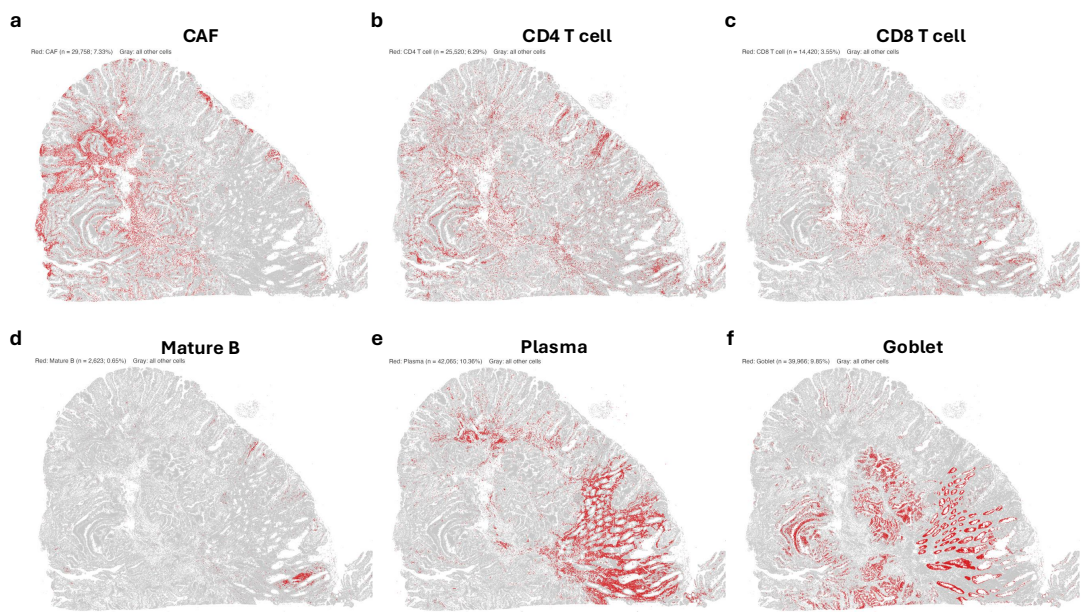

Fig. S14: Individual spatial map for normal cells labeled by Seurat in Xenium

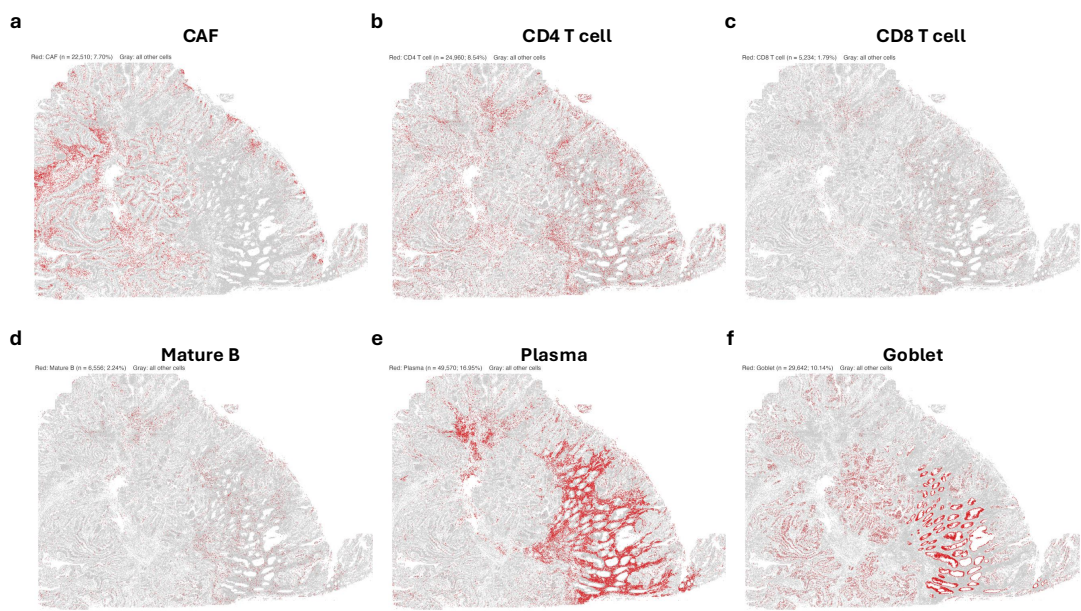

Fig. S15: Individual spatial map for normal cells labeled by Seurat in CosMx

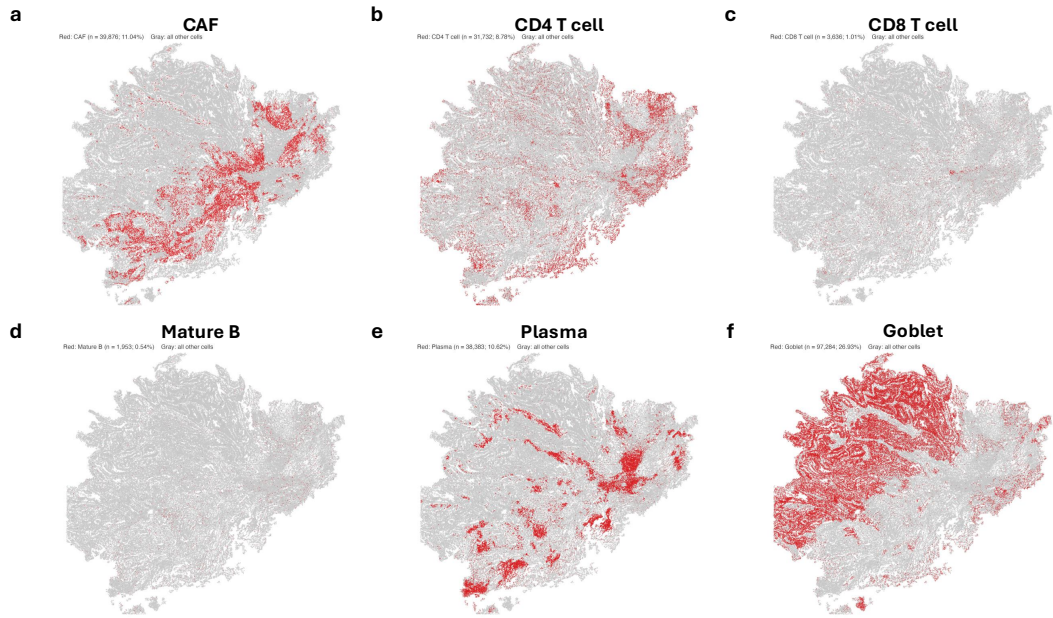

Fig. S16: Individual spatial map for normal cells labeled by Seurat in Stereo-seq

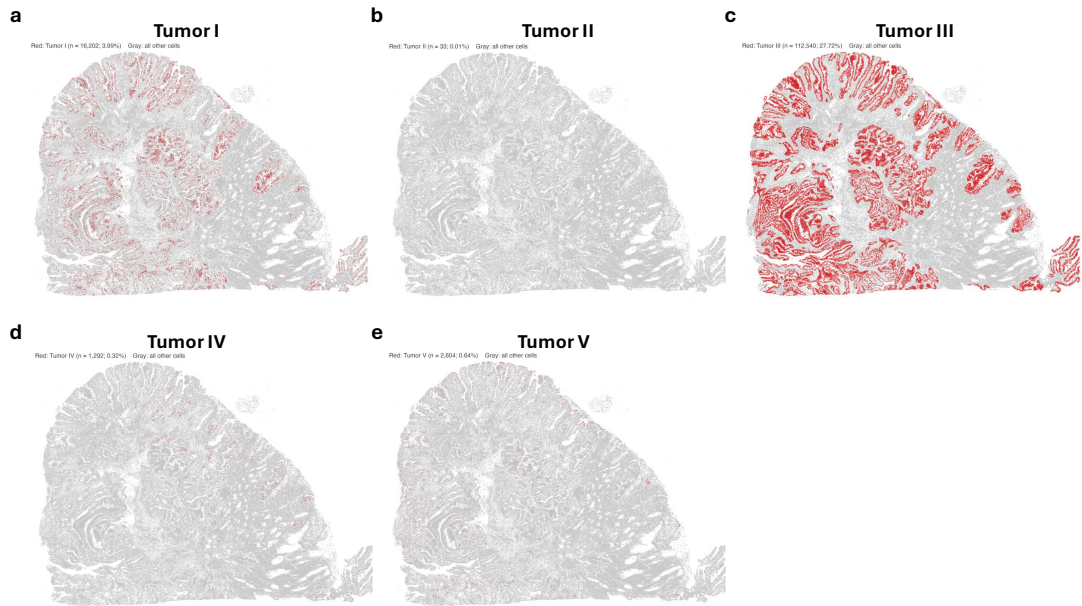

Fig. S17: Individual spatial map for tumor cells labeled by Seurat in Xenium

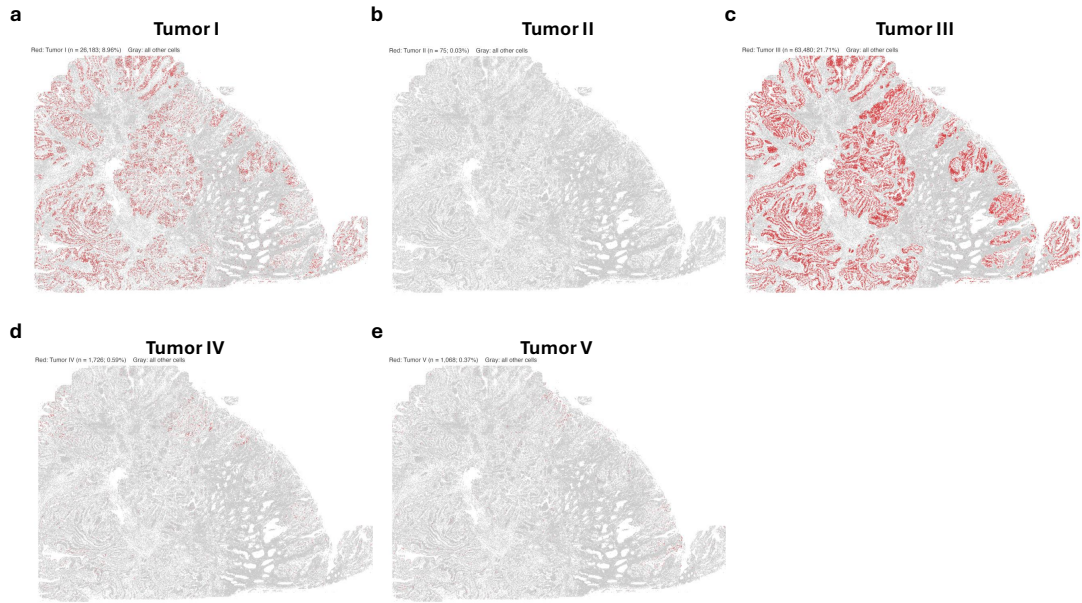

Fig. S18: Individual spatial map for tumor cells labeled by Seurat in CosMx

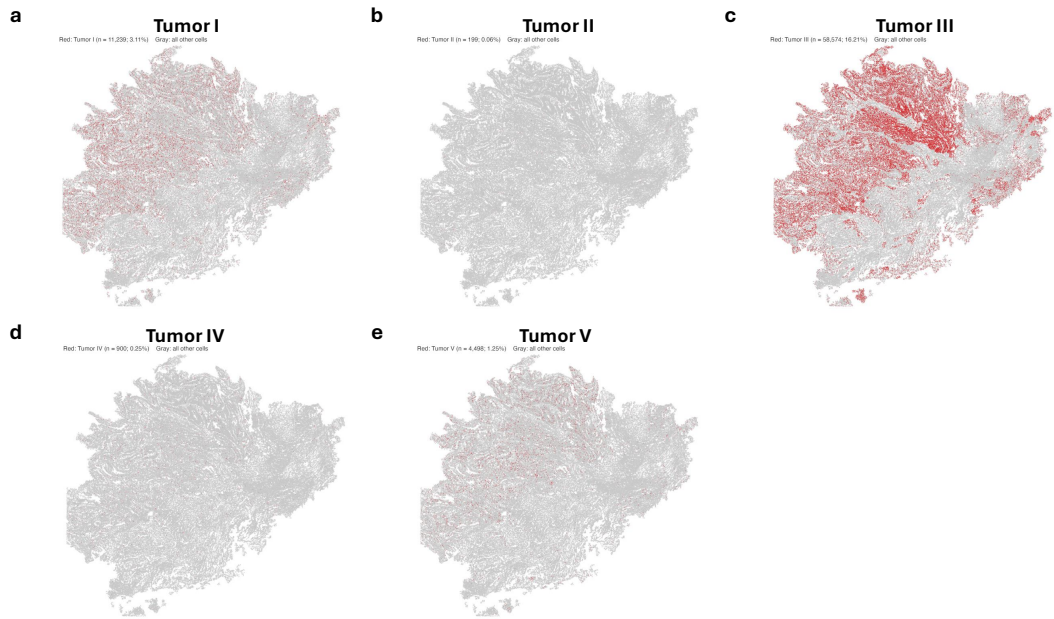

Fig. S19: Individual spatial map for tumor cells labeled by Seurat in Stereo-seq

##### 3.1.3 SingleR

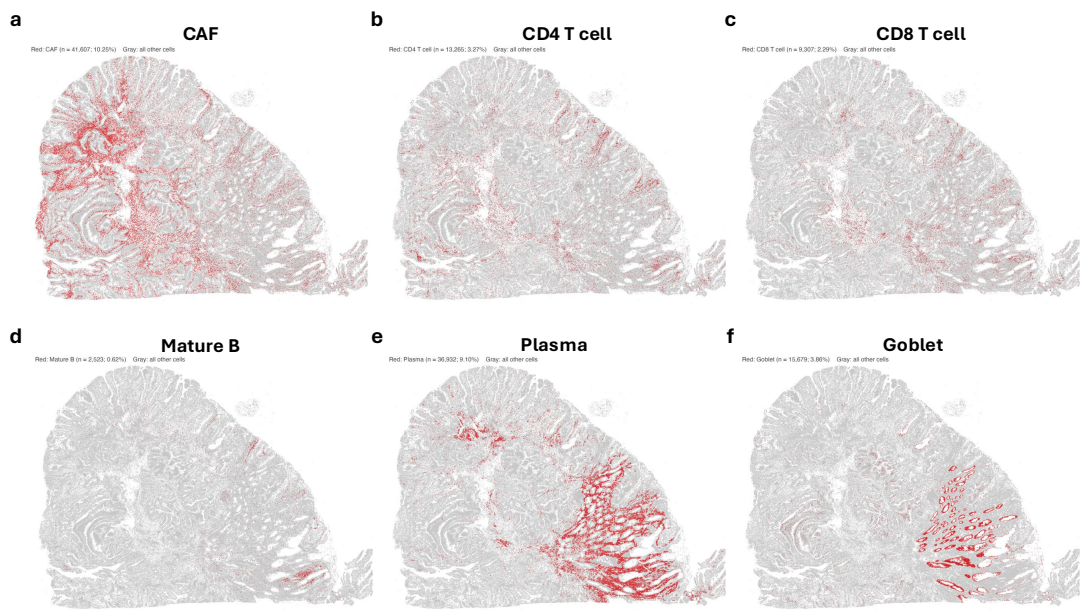

Fig. S20: Individual spatial map for normal cells labeled by SingleR in Xenium

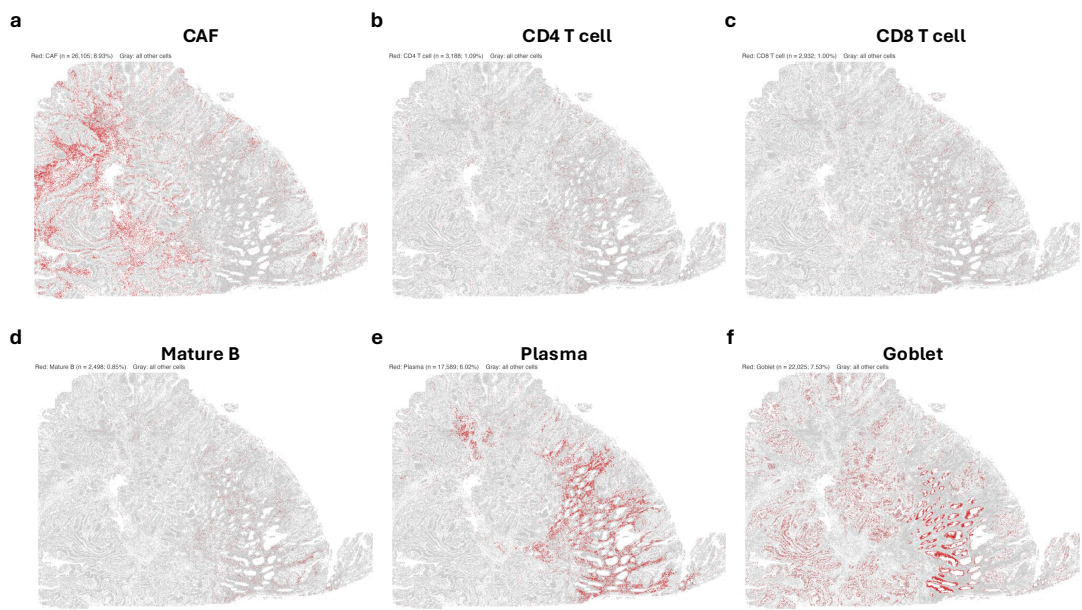

Fig. S21: Individual spatial map for normal cells labeled by SingleR in CosMx

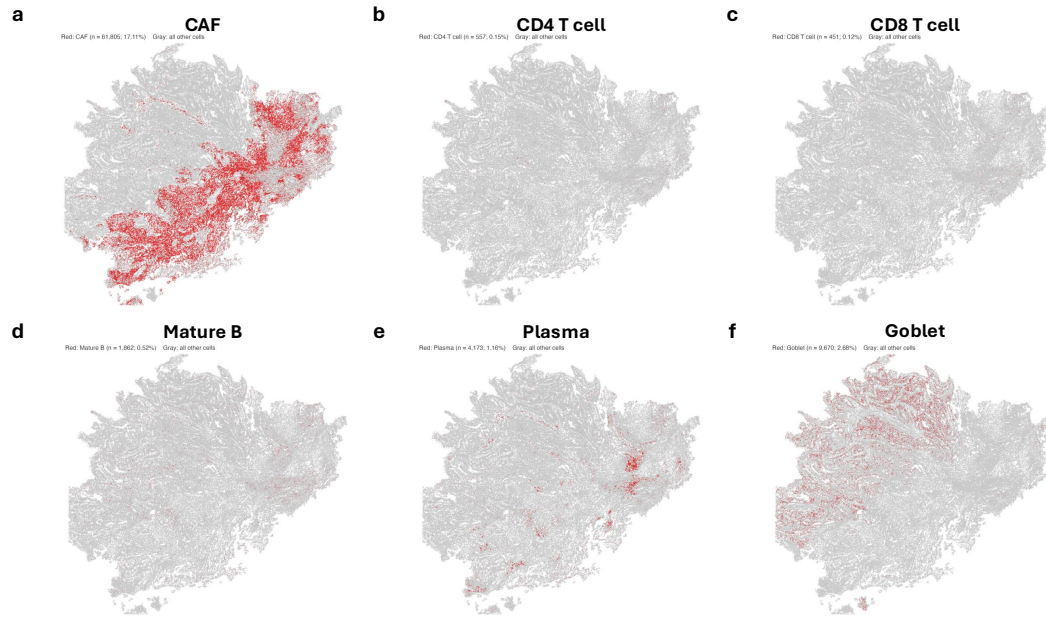

Fig. S22: Individual spatial map for normal cells labeled by SingleR in Stereo-seq

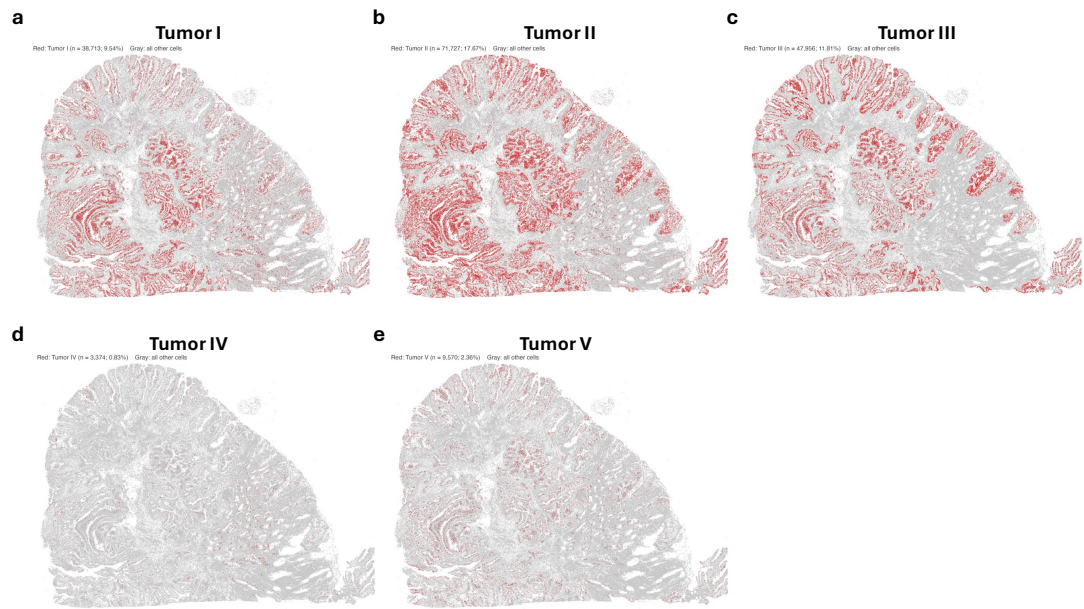

Fig. S23: Individual spatial map for tumor cells labeled by SingleR in Xenium

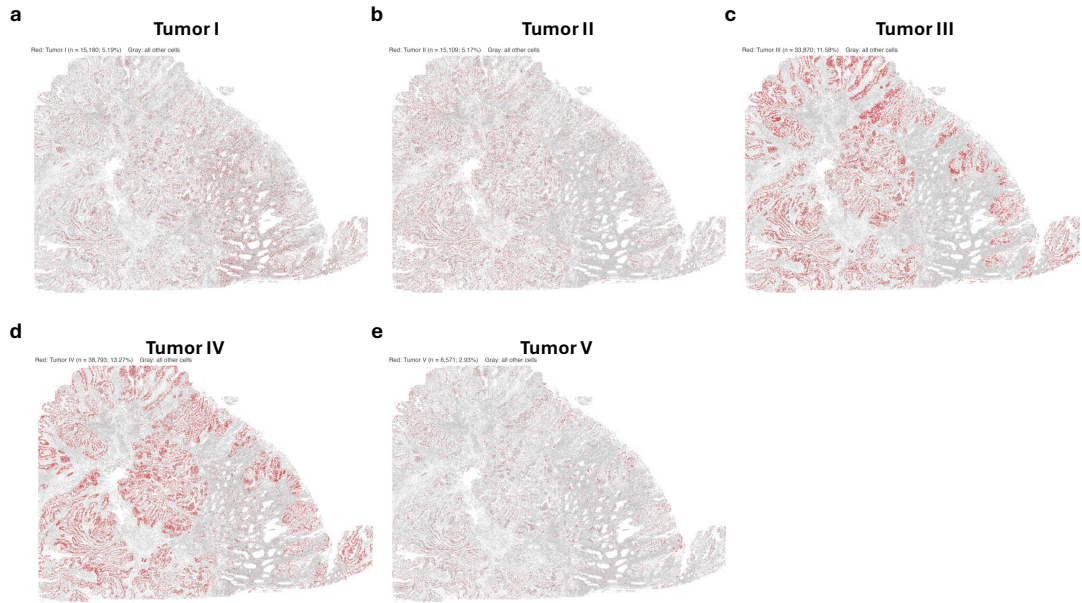

Fig. S24: Individual spatial map for tumor cells labeled by SingleR in CosMx

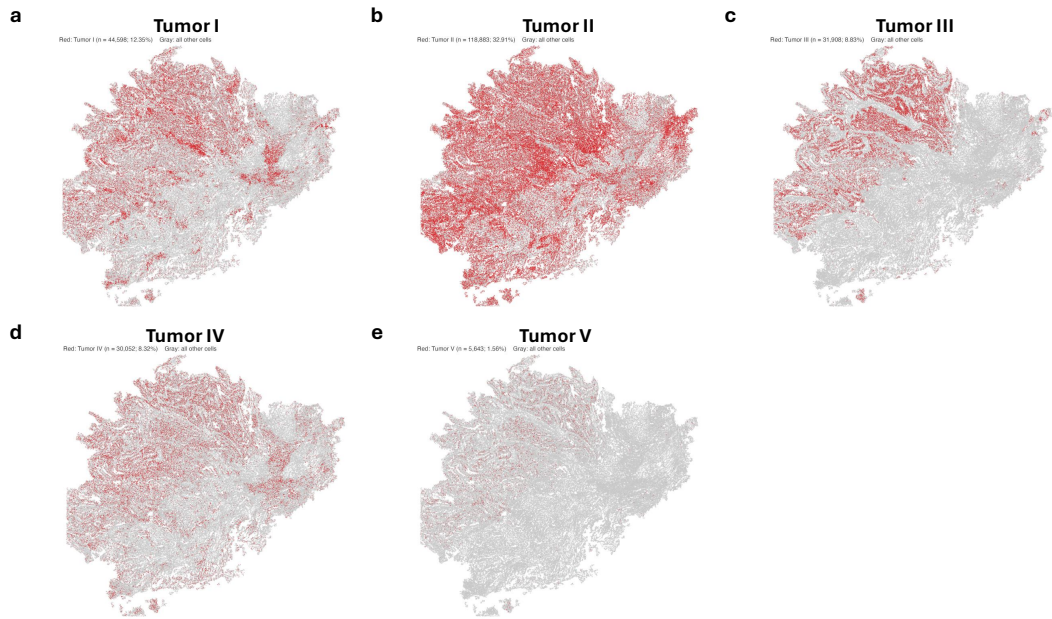

Fig. S25: Individual spatial map for tumor cells labeled by SingleR in Stereo-seq

##### 3.1.4 CellTypist

The cross-platform results by CellTypist are visualized below, showing poor performance in tumor cell annotation and in CosMx compared to BiNVaST, Seurat, SingleR.

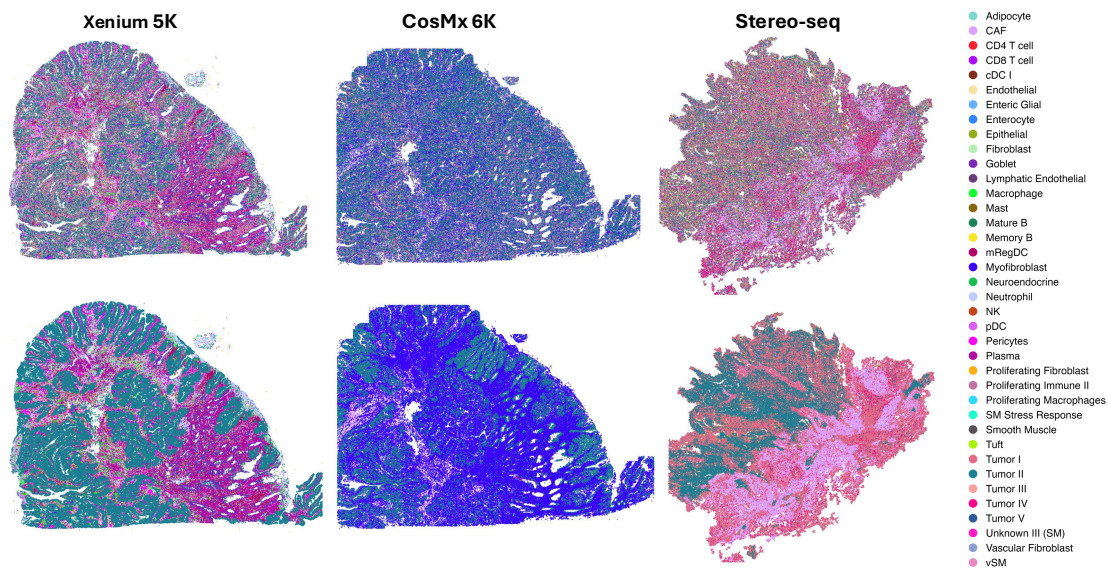

Fig. S26: Spatial maps for cell labels predicted by CellTypist across three platforms. The first row is raw predicted cell label and the second is majority-voted labels.

#### 4 Pediatric Diffuse Hemispheric Glioma

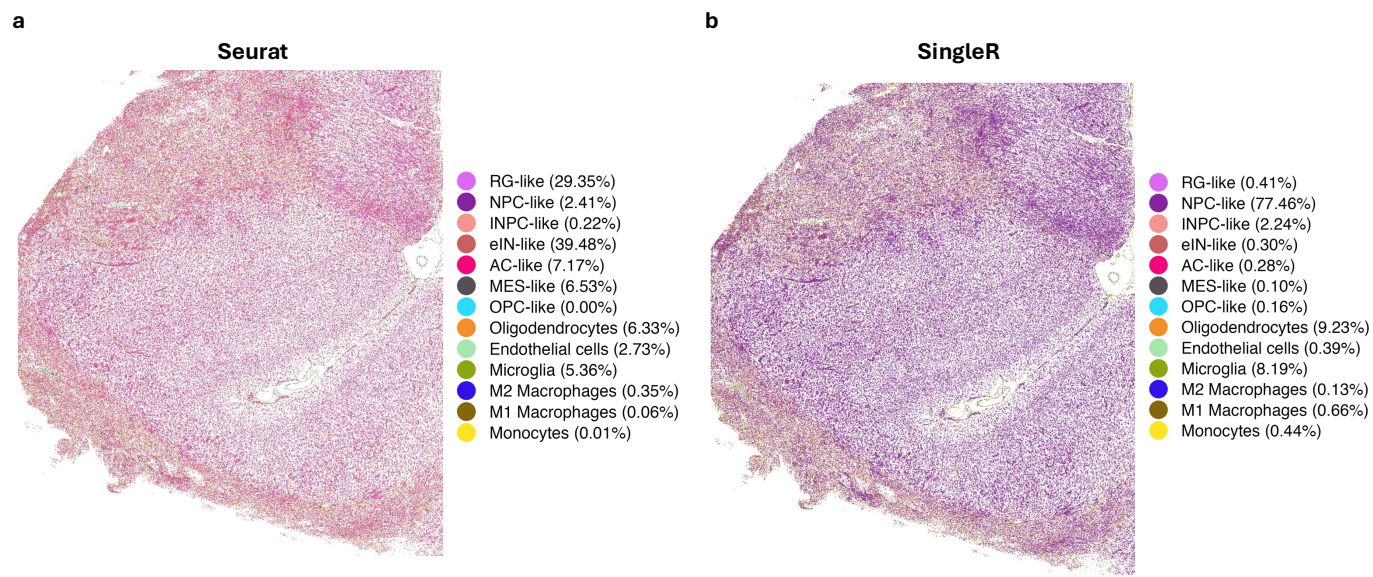

Fig. S27: Spatial maps for cell labels in DHG tumor predicted by Seurat and SingleR.
